# A fibroblast-CD8+ T cell circuit drives a Type 1 shift in allergic inflammatory memory

**DOI:** 10.64898/2026.11.02.763646

**Authors:** Andrew C Kwong, Isabelle Oliver, Lixin Yang, Peter Lotfy, Jose Ordovas-Montanes

## Abstract

Inflammatory memory in barrier tissues often manifests as exaggerated recall responses to previously encountered stimuli ^1,2^. While recall is generally thought to reinforce the Type of inflammation (e.g., Type 1 (T1), Type 2 (T2), or Type 17 (T17)) induced in the primary encounter, whether and how memory responses engage programs beyond the initiating Type is unclear. Here, we adapted well-characterized murine models of T2 atopic dermatitis (AD)-like ^3^ and T17 psoriasis-like ^4^ inflammation into repeated challenge paradigms to compare how inflammatory memory is stored and recalled through longitudinal single-cell RNA-sequencing. In T17 inflammation, a network of IL-17-producing γδT cells, IL-17-responsive keratinocytes and myeloid cells established in primary challenge re-emerged in recall with increased abundance and accelerated kinetics. By contrast, allergic T2 memory drove a particularly exacerbated and durable recall response in which hallmark features of T1 immunity, including interferon (IFN) signaling and cytotoxic lymphocyte recruitment, were unexpectedly layered onto amplified T2 responses. This T2-to-T1 shift was initiated by CD8+ tissue-resident memory T (Trm) cells that expanded with repeated challenge, produced IFNγ, TNF-α and IL-13, and closely resembled treatment-refractory Trm cells in AD. Tissue rewiring was supported by Trm-derived T1 cytokines and the IL-4-rich T2 milieu whose synergy on fibroblasts amplified allergic recall via IL-15 and *CCL19*. By tuning fibroblast IFN responsiveness in vivo, we mechanistically dissect how this fibroblast-CD8+ T cell circuit supports cytotoxic CD8+ T cell responses. In human AD, both arms of allergic recall amplification by this two-cell circuit featuring T1 and T2 effector responses were conserved. Our findings reveal how a stromal-immune circuit drives a clinically relevant T1 immune axis to support allergic inflammatory memory.

## Introduction

The ability of tissues to remember prior inflammatory experiences, broadly termed inflammatory memory, is a fundamental property of host defense. Cells and tissues retain information from prior encounters through changes in epigenetic state, cellular identity and tissue organization that collectively reshape responses to future stimuli ^1,2^. These adaptations underlie diverse biological processes ranging from enhanced protection against recurrent infection to accelerated wound healing after injury ^5,6^. However, the same mechanisms can become maladaptive and promote exaggerated immune responses that drive disease^7^. Defining how inflammatory memory is distributed and re-engaged in tissues is therefore central to advancing our understanding of both protective immunity and chronic inflammatory disease.

Tissue inflammatory responses can be categorized into three major classes of immunity that coordinate specialized arms of host defense–Type 1 (T1) cytotoxic, Type 2 (T2) allergic and Type 17 (T17) antifungal–each defined by distinct instructive and effector cytokine programs ^8^. In many settings, inflammatory memory manifests as a Type-specific exaggeration in which repeated challenge reinforces the immune program established by the initial response ^9^. In T1 immunity, primary viral infection induces an interferon (IFN) response that recruits immune effectors in waves over two weeks ^10^. Nonrecirculating tissue-resident memory T (Trm) cells seeded after viral clearance rapidly produce IFNγ upon rechallenge to recruit these same effector populations within two days^10,11^. In the small intestine, T2 responses to primary parasite infection expand group 2 innate lymphoid cells (ILC2s)^12^, enhance their production of effector cytokines ^13^, and induce T helper 2 (Th2) cell differentiation ^14^. Together, these changes in tissue composition and cell states encode a memory that protects against re-infection ^14,15^.

Type-specific memory responses, however, can also stabilize chronic inflammatory disease. In psoriasis–a T17 inflammatory skin disorder–IL-17-producing CD8+ Trm (Trm17) cells persist in clinically resolved lesions ^16^. The activation of T cells in explants of resolved skin is sufficient to induce T17 epithelial responses whose strength correlates with poor treatment responsiveness ^17,18^. Skin-resident dendritic cells (DCs) also retain memory states that enhance IL-23 production ^19^, reinforcing T17-specific exaggerations to drive disease. Similarly, in a murine model of allergic asthma, memory is stored in lung-resident allergen-specific CD4+ Trm cells^20,21^. Allergen re-exposure activates these Trm cells, which rapidly initiate T2 responses that recruit eosinophils and Th2 cells responsible for pathology ^20,21^. Collectively, these studies support the prevailing paradigm in which both protective and maladaptive inflammatory memories reinforce the inflammatory Type associated with the primary response.

Nevertheless, many of these insights have emerged from targeted interrogation of the inflammation Types expected in primary responses. Unbiased high-resolution approaches could thus stand to reveal a more complex interplay of inflammatory programs across memory encoding and recall.

Emerging work has broadened the concept of tissue memory beyond its storage in tissue-resident immune cells to encompass its distribution and recall across multicellular ecosystems. Landmark studies have shown that epithelial and stromal cells can themselves encode durable inflammatory memories through epigenetic remodeling ^5,22,23^. For instance, epithelial stem cells in the respiratory epithelium of chronic rhinosinusitis patients retain an intrinsic memory that enhances sensitivity to IL-4 and IL-13^23^. Simultaneously, memory recall–defined by the inflammatory circuitry engaged in secondary challenge–is increasingly recognized to involve coordinated interactions between immune and structural cells. In the T1 skin disease vitiligo, IFNγ from autoreactive T cells induces keratinocyteand fibroblast-derived chemokines that recruit CD8+ T cells to drive a positive feedback loop for melanocyte destruction ^24,25^. Separately, small intestinal helminth infection reveals how memory can shift cellular dependencies in inflammation. While epithelial tuft cells are dispensable for priming Th2 cells in primary infection, they are essential for pathogen clearance in rechallenge by activating memory Th2 cells^14^. We are only beginning to understand the principles through which cell circuits organize the encoding and recall of memory responses across Types of inflammation.

To address this knowledge gap, we turned to two wellestablished murine models of sterile skin inflammation with highly stereotyped Type-specific immune responses: MC903-induced T2 inflammation and imiquimod (IMQ)-induced T17 inflammation ^3,4^. MC903 models atopic dermatitis (AD) by inducing keratinocyte TSLP production ^3^. This alarmin initiates a T2 inflammatory cascade that promotes the activation of basophils and Th2 cells, which drive pathology via IL-4 and IL-13^26,27^. Modeling T17 responses, IMQ induces psoriasis-like inflammation through IL-23 from DCs, which in turn drives IL-17 from dermal γδT cells^4,28^. Repeated challenge, in which acute bouts of IMQ are separated by clinical resolution, is an established model of T17 inflammatory memory. Individual memory states are encoded in IL-17-producing γδT cells and epithelial stem cells^5,29,30^.

By contrast, few studies have characterized T2 inflammatory memory in skin ^31–33^. Through an unbiased comparison of T2 and T17 responses in skin, we sought to define how Type-specific memory encodings re-organize the tissue circuitry engaged upon recall.

To study the organizing principles of inflammatory memory, we adapted acute T2 and T17 models of skin inflammation into repeated challenge paradigms. Longitudinal profiling of skin ecosystem composition by ingle-cell RNA-sequencing (scRNA-seq) revealed that T17 memory conformed to the prevailing paradigm of Type-specific exaggeration. A similar network involving keratinocytes, γδT cells and neutrophils emerged across primary and secondary T17 responses; recall featured increased abundances of these subsets appearing with accelerated kinetics. Relative to T17 memory, allergic memory involved a particularly exacerbated response that was durable and specific to T2 rechallenge. Recall not only amplified hallmark features of T2 inflammation but also unexpectedly acquired characteristics of T1 immunity. Allergic CD8+ Trm cells initiated this T2-to-T1 shift, which was then amplified by a fibroblast-CD8+ T cell circuit whose features were conserved in AD and refractory to treatment. Altogether, we demonstrate how the engagement of an immune-stromal cell circuit motif upon allergic memory recall rewires circuitry in the skin to promote a T1 shift that is central to AD pathology.

## Results

### Repeated T2 challenge encodes durable, site-specific allergic inflammatory memory

To compare how inflammatory memory is stored and recalled across distinct inflammatory Types, we established dual-challenge models of T2 and T17 skin inflammation. For T17 inflammation, we utilized the previously established IMQ recall protocol, in which prior IMQ challenge enhances secondary responses through γδT cells and changes in keratinocyte chromatin accessibility ^5,29,30^. In parallel, we adapted this setup for MC903 to assess whether T2 inflammation similarly establishes persistent tissue memory ^3^. For each inflammation Type, murine ear skin received a six-day primary challenge (D0–D6), a 30-day recovery period, and a six-day secondary challenge (D36–D42) (Fig 1A). As expected, mice previously exposed to IMQ exhibited approximately two-fold greater ear thickening during secondary challenge (Fig 1B). Hematoxylin and eosin (H&E) staining confirmed greater epidermal hyperplasia and immune infiltration in T17 memory recall than in primary T17 inflammation (Fig 1C).

**Figure 1.**
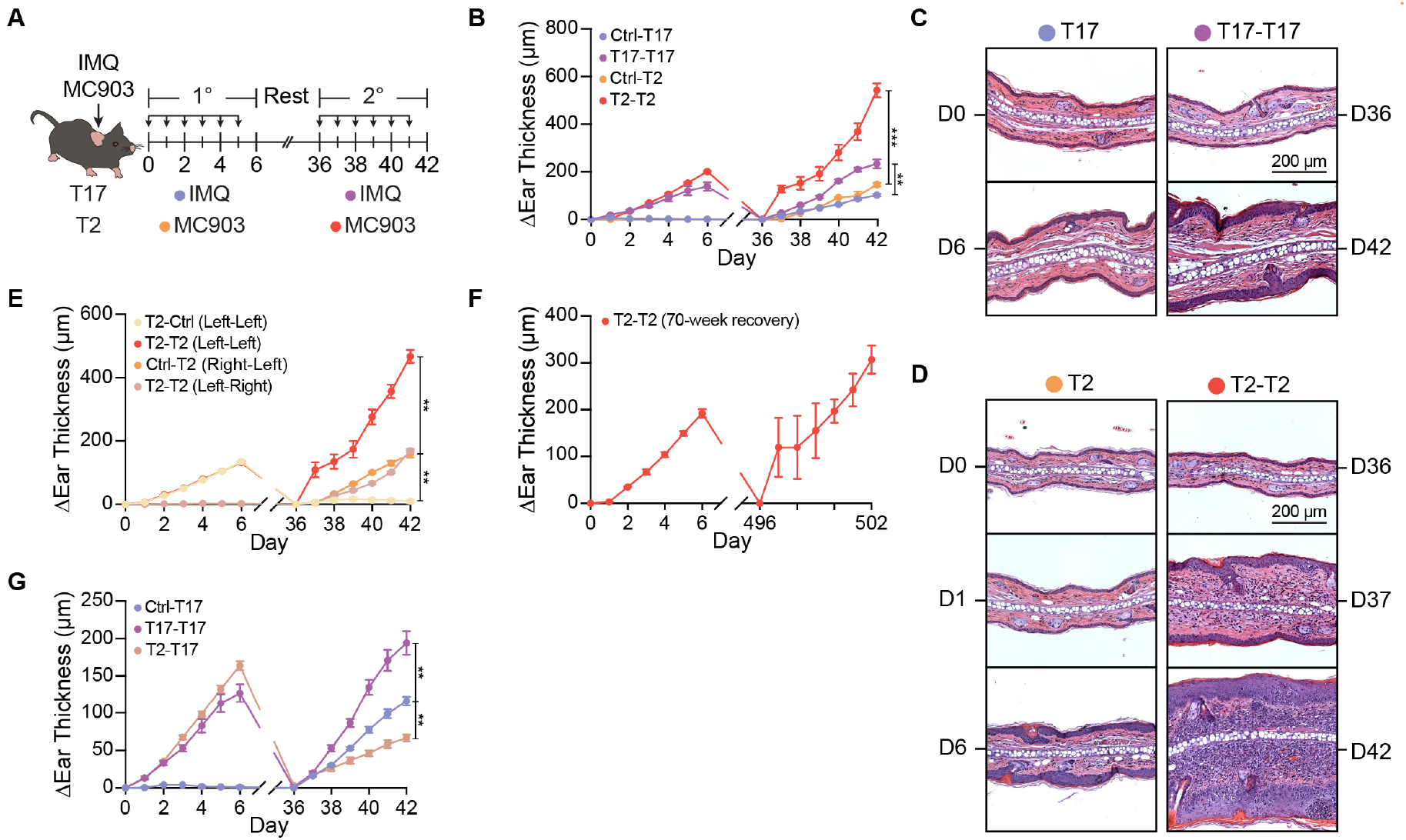
Allergic inflammatory memory is a rapidly exaggerated, site-specific, durable and Type-specific response. **(A)** Schematic of repeated challenge inflammatory models of T17 and T2 inflammation. **(B)** Ear thickness curves across primary and secondary inflammatory challenges (n=5 mice/group). Representative H&E images for acute and memory T17 inflammation at baseline and end of six-day treatment period. **(D)** Representative H&E images of baseline, one-day and six-day ear skin for acute and memory T2 inflammation. **(E)** Ear thickness curves of site-matched (left-left) or unmatched (left-right) T2 memory (n=5 mice/group). **(F)** Ear thickness curves of allergic memory with long-term (>1Y) rest (n=3 mice). **(G)** Ear thickness curves of T17 rechallenge after T2 primary challenge (n=12 (Ctrl-T17), 7 (T17-T17), 15 (T2-T17) mice).

Strikingly, prior MC903 exposure led to a markedly stronger response at secondary challenge than T17 memory. Six days of recall (D42) produced four-fold greater ear thickening than six days of primary challenge (D6) (Fig 1B). This response emerged rapidly: ear thickness in one day of recall (D37) approached the six-day peak of primary inflammation (Fig 1B). H&E staining indicated that this early response primarily reflected immune infiltration rather than edema and was further exacerbated with continued treatment (Fig 1D). Importantly, the 30-day recovery period permitted histological normalization before rechallenge (Fig 1C–D). Although a modest residual increase in ear thickness was observed after T2 recovery (D36), trichrome staining showed no clear evidence of fibrosis (Extended Data Fig 1A–B). Thus, while both inflammatory Types displayed enhanced secondary responses, T2 memory was distinguished by its particularly rapid and exaggerated recall phase.

The magnitude of the T2 recall response relative to T17 memory prompted us to better define the properties of allergic inflammatory memory. First, we examined how primary challenge duration and stimulus dose influenced memory. At least five days of primary MC903 treatment were required to generate a memory response upon rechallenge (Extended Data Fig 1C). We also found that halving the primary dose of MC903 did not alter recall, in contrast to the clear dose dependence of T17 memory (Extended Data Fig 1D–E). We then asked whether recall reflected a localized tissue memory or a systemic change in inflammatory responsiveness. Critically, secondary MC903 produced an enhanced response only when applied to T2-experienced skin and not to the contralateral vehicle-treated ear (Fig 1E). Circulating factors alone are thus not sufficient to produce T2 memory, which remains anatomically restricted to the site of primary inflammation.

To assess the durability of T2 memory, we extended the recovery period after primary challenge to 70 weeks. Remarkably, the exaggerated one-day ear thickening response and associated histopathology remained intact despite this prolonged recovery interval (Fig 1F, Extended Data Fig 1F). Finally, we asked whether T2 memory broadly enhanced subsequent inflammatory responses or instead remained selective for the initiating inflammation Type. Interestingly, replacing MC903 with IMQ during secondary challenge suppressed the T17 inflammatory response (Fig 1G). Together, these findings establish a dual-challenge model in which T2 inflammation gives rise to site-specific, durable and Type-selective tissue memory.

### Allergic inflammatory memory produces a T1 shift in skin ecosystem composition

To examine how skin tissue ecosystems are remodeled during T2 and T17 inflammatory memory, we conducted longitudinal scRNA-seq. As differences between primary and recall responses emerged within three days of treatment (Fig 1B), we selected timepoints spanning this window in primary (D0-4, D6) and secondary (D36-39) challenge, hashing 3 biological replicates per timepoint (Fig 2A). After doublet removal and quality control filtering, we recovered 246,250 single-cell transcriptomes from 60 mice across primary and secondary T2 and T17 inflammation (Fig 2A; Methods).

**Figure 2.**
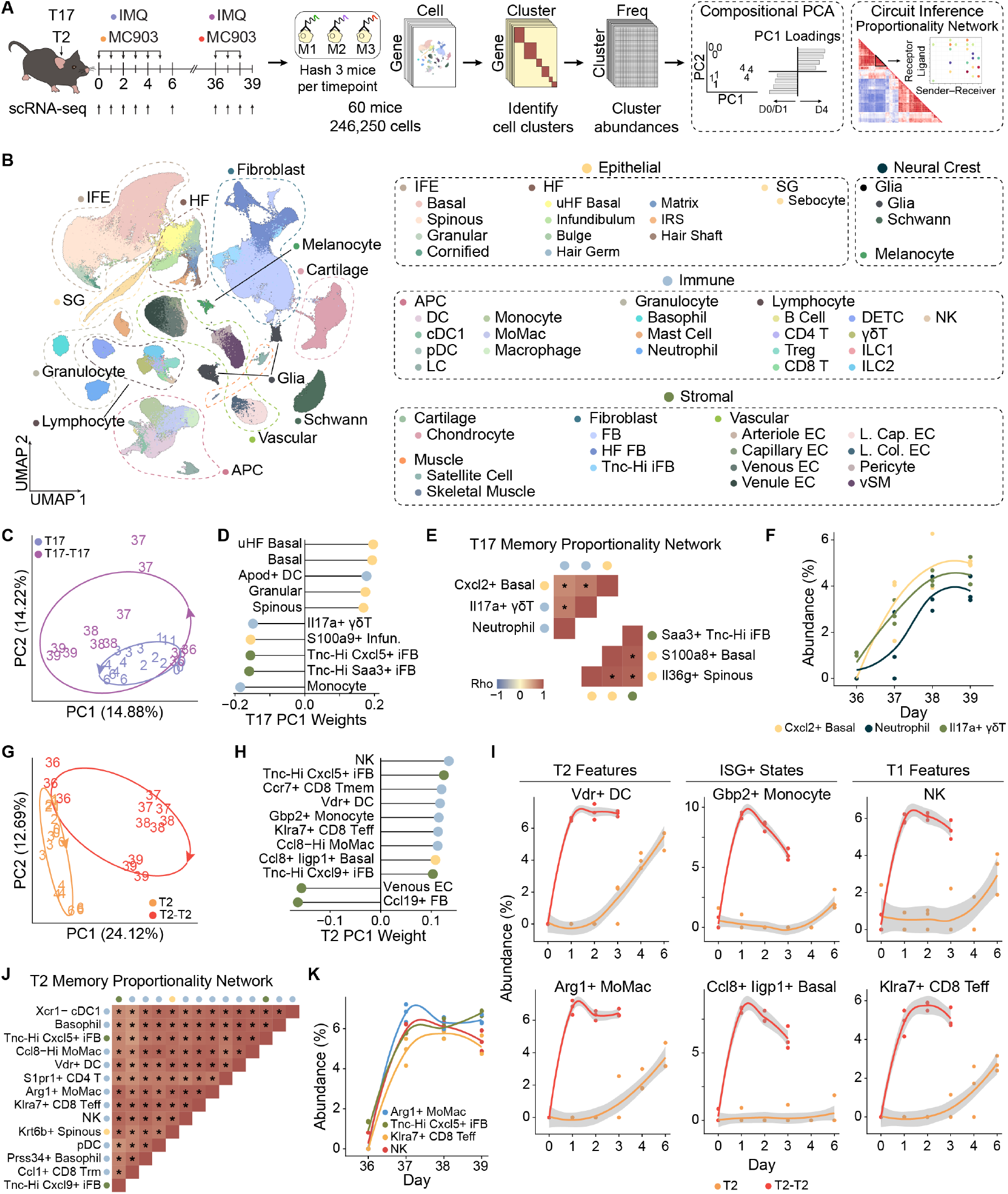
Compositional analysis of allergic skin memory reveals a T2-to-T1 shift in recall. **(A)** Schematic of compositional analysis workflow across T17 and T2 inflammation with three mice hashed per timepoint. Cells are annotated into clusters by iterative clustering, which are then used to generate a cluster abundance matrix for each mouse. Compositional PCA is used to identify individual cell clusters that drive longitudinal changes while proportionality analysis infers multicellular networks of co-occurring clusters. **(B)** Uniform manifold approximation and projection (UMAP) embedding of 246,250 cells across 60 mice coloured by cell type and subtype with markers highlighted in Extended Data Fig 2A-D. **(C)** Compositional PCA of T17 inflammation samples where each point is annotated by timepoint and coloured by timecourse. Ellipse is centered on the centroid for each timecourse. Distance reflects variation in cell cluster abundance on a per-sample basis with PC1 cluster abundance loadings. **(E)** Representative significant cell networks taken from proportionality heatmap calculated across T17 memory timepoints with full network shown in Extended Data Fig 4. A significance cutoff FDR < 0.05 generated by permutation testing was used to identify significant relationships between cell clusters. **(F)** Abundance plot of *Cxcl2*+ basal keratinocytes, neutrophils and *Il17a*+ γδT cells in T17 memory. **(G)** Compositional PCA of T2 inflammation samples with **(H)** PC1 loadings. **(I)** Abundance plots across primary and secondary T2 inflammation for features nominated from PC1. **(J)** Representative significant cell network calculated across T2 memory timepoints with full network shown in Extended Data Fig 4. **(K)** Co-occurrence of *Arg1*+ monocytederived macrophages (MoMac), NK cells, *Klra7*+ CD8 effector T (Teff) cells and *Cxcl5*+ *Tnc*-Hi inflammatory fibroblasts (iFBs) are plotted as each cluster’s abundance within the total skin ecosystem.

Unsupervised top-level clustering separated 4 major cellular categories: epithelial, immune, stromal and neural crest (Fig 2B). We then performed subclustering based on canonical marker genes to identify 48 cell types such as basal keratinocytes, DCs and fibroblasts (Fig 2B, Extended Data Fig 2A-D). Through iterative clustering, we identified 150 distinct end cell clusters that captured the diversity of cell types, subsets and states in murine skin (Supplementary Table 1). Among epithelial cells, *Tslp*+ spinous keratinocytes were induced by MC903, specific to T2 inflammation and coexpressed *Cyp24a1*, a vitamin D-metabolizing enzyme that may negatively regulate MC903 responses (Extended Data Fig 2E-F)^3,34^. Meanwhile, *Il36g*+ spinous keratinocytes were enriched during T17 inflammation and expressed psoriasisassociated IL-17-induced antimicrobial peptides including *S100a8* and *Sprr2d* (Extended Data Fig 2E-F)^35,36^. Targeted examination of keratinocyte clusters thus establishes that MC903 and IMQ recapitulate core features of epithelial responses across T2 and T17 skin inflammation.

Beyond epithelial cells, fibroblasts have also emerged as key organizers of immune responses in skin during T2 and T17 inflammation. We recovered 43,506 fibroblasts comprising 15 transcriptionally distinct clusters, including three inflammatory fibroblast (iFB) populations marked by the extracellular matrix protein *Tnc* (Extended Data Fig 2G). Consistent with prior work in psoriasis^37,38^, *Saa3*+ *Tnc*-Hi iFBs were enriched in T17 inflammation (Extended Data Fig 2H). While this state was also present throughout MC903 treatment, *Tnc*-Hi iFB states marked by *Cxcl5*+ and *Cxcl9*+ were more abundant in T2 inflammation (Extended Data Fig 2H). Recent work has shown that *Tnc*+ iFBs promote acute T2 responses to MC903 through IL-4Rα signaling ^27^. Our atlas therefore recapitulates iFB populations described across allergic and psoriasiform skin inflammation, providing a foundation to examine their potential contributions to memory responses.

To unbiasedly identify drivers of inflammatory memory, we treated each skin sample as an ecosystem in which individual cell clusters represent “species” whose abundances changed over time ^10,39^. Within each mouse replicate, we quantified the proportional representation of each cluster relative to all cells in that sample to generate a cluster abundance matrix (Fig 2A). This matrix was then used for two complementary analyses. First, we performed principal component analysis (PCA) to identify individual cell clusters whose abundances drove temporal variation in memory (Fig 2A). Second, we conducted proportionality-based co-occurrence analysis to infer coordinated population dynamics between cell clusters (Fig 2A) ^10,40^. By applying these analyses to our cell state abundances, we identified groups of clusters with co-occurring longitudinal dynamics, which we term inferred multicellular networks. The relationships between individual cluster nodes could then be further characterized by cell-cell communication inference, followed by experimental validation to formally define cell circuits. Overall, this framework leverages the temporal resolution of our sampling strategy to nominate multicellular networks and candidate cell circuits contributing to inflammatory memory.

To test our compositional framework, we first asked whether it could recover established features of T17 inflammatory memory. PCA revealed that late primary T17 inflammation (D4-6) separated from early timepoints along the negative PC1 axis, with loadings that included *Il17a*+ γδT cells, *S100a9*+ infundibulum keratinocytes and monocytes (Fig 2CD). Together, these populations captured key aspects of T17 psoriasiform inflammation: IL-17-producing T cells, epithelial antimicrobial peptide secretion and myeloid cell recruitment ^36,41^. Notably, T17 secondary challenge represented an accelerated and exaggerated version of this initial inflammatory program, occupying a similar compositional space as primary challenge (Fig 2C). After only four days of secondary challenge, the D39 timepoint aligned with the D6 endpoint of primary T17 inflammation (Fig 2C, Extended Data Fig 3A). Similarly, PC2 captured early IFN-stimulated cell states that emerged one day after both primary and secondary challenge but exhibited accelerated kinetics during recall (Extended Data Fig 3B-C), consistent with established roles for type I IFN in psoriasis initiation ^42^. Thus, T17 memory is marked by accelerated and exaggerated re-emergence of the cell clusters that drive the primary response, effectively reinforcing the original inflammatory ecosystem.

To examine coordinated cell state dynamics in T17 inflammation, we used proportionality analysis to reconstruct a cooccurrence network with 56 significant relationships between cell clusters out of 11,250 possible pairs (Extended Data Fig 4). We identified a putative network comprising *Il17a*+ γδT cells, *Cxcl2*+ basal keratinocytes and neutrophils that displayed coordinated abundance dynamics in T17 memory (Fig 2E-F). Notably, these core interactions closely mirror prevailing models of psoriasis pathogenesis in which IL-17-producing lymphocytes stimulate keratinocytes to recruit neutrophils ^36,41^. Our co-occurrence network also linked *Il36g*+ spinous keratinocytes and *Saa3*+ *Tnc*-Hi iFBs, two populations that have been independently implicated in IMQ pathogenesis but not previously connected within the same inflammatory network (Fig 2E)^37^. Altogether, our compositional framework both recovers established inflammatory networks and generates novel hypotheses for cell circuits that may support memory establishment and recall.

Having observed that T17 memory involves a reinforcement of the primary inflammatory response, we next asked whether T2 inflammatory memory behaved similarly. PCA revealed that D4-D6 of acute T2 inflammation separated along the negative PC2 axis (Fig 2G). Loadings included expected populations such as *Tslp*+ spinous keratinocytes, the key state induced by MC903 ^3^, together with canonical AD-associated effectors like *Mki67*+ CD4 T cells, *Mki67*+ ILC2s and Tregs (Extended Data Fig 3D-E)^43,44^. Interestingly, *Cxcl5*+ *Tnc*-Hi iFBs arose as one of the strongest drivers of PC2 (Extended Data Fig 3D-E). This unbiased observation aligns with prior work showing that IL-4-responsive *Tnc*+ fibroblasts recruit immune cells during T2 skin inflammation ^27,45^. Notably, analogous *TNC*+ fibroblast populations have also been identified in human AD skin^46^.

In contrast to T17 memory, T2 recall occupied a distinct compositional space that diverged from the primary inflammatory response (Fig 2G). Rather than simply exaggerating acute T2 inflammation along the PC2 axis, secondary challenge introduced novel memory-specific tissue ecosystem states. Canonical T2-associated populations, including *Ccl8*-Hi monocyte-derived macrophages and *Vdr*+ DCs^44,47^, displayed accelerated and exaggerated emergence kinetics relative to primary challenge, mirroring patterns from T17 memory (Fig 2H-I). Unexpectedly, recall-specific cell states were enriched for canonical features of T1 immunity. IFNstimulated populations like *Ccl8*+ *Iigp1*+ basal keratinocytes and *Cxcl9*+ *Tnc*-Hi iFBs were uniquely induced during secondary challenge (Fig 2H-I). In parallel, we observed rapid recruitment of T1 immune populations including *Klra7*+ CD8+ effector T (Teff) cells and NK cells (Fig 2H). Thus, T2 inflammatory memory is characterized not only by exaggerated T2-associated responses but also by the acquisition of hallmark T1 immune features during recall.

Proportionality analysis identified 2,265 significant relationships between clusters constituting approximately 20% of possible pairs (Extended Data Fig 4). The T2 co-occurrence network was therefore markedly more interconnected than the inferred T17 network (Extended Data Fig 4). Strikingly, T1 immune populations like pDCs and NK cells co-occurred with canonical T2 clusters such as basophils and *Arg1*+ monocyte-derived macrophages, suggesting their concurrent emergence in allergic recall (Fig 2J-K). Furthermore, *Cxcl5*+ *Tnc*-Hi and *Cxcl9*+ *Tnc*-Hi iFBs appeared in this inferred network alongside mixed T1 and T2 features. The amplifying role of iFBs in acute T2 inflammation may thus extend into recall. Altogether, our unbiased approach reveals how allergic memory encodes a recall-specific T2-to-T1 shift in contrast to the Type-specific exaggeration of T17 memory.

### CD8+ Trm cells drive the T1 shift in allergic inflammatory memory

(A)To identify the drivers of this T1 shift, we examined the particularly exacerbated one-day response of T2 recall. We generated pseudobulk profiles for each biological sample on the first day of primary (D1) and secondary (D37) challenge for differential expression analysis. Gene set enrichment analysis (GSEA) nominated the IFNγ response as the most enriched pathway on D37, followed by IFNα and inflammatory response pathways (Fig 3A). ELISA measurements of skin lysates confirmed robust IFNγ induction on D37 that was absent in primary challenge (Fig 3B). By contrast, IFNα and IFNβ were not significantly induced during recall (Extended Data Fig 5A).

**Figure 3.**
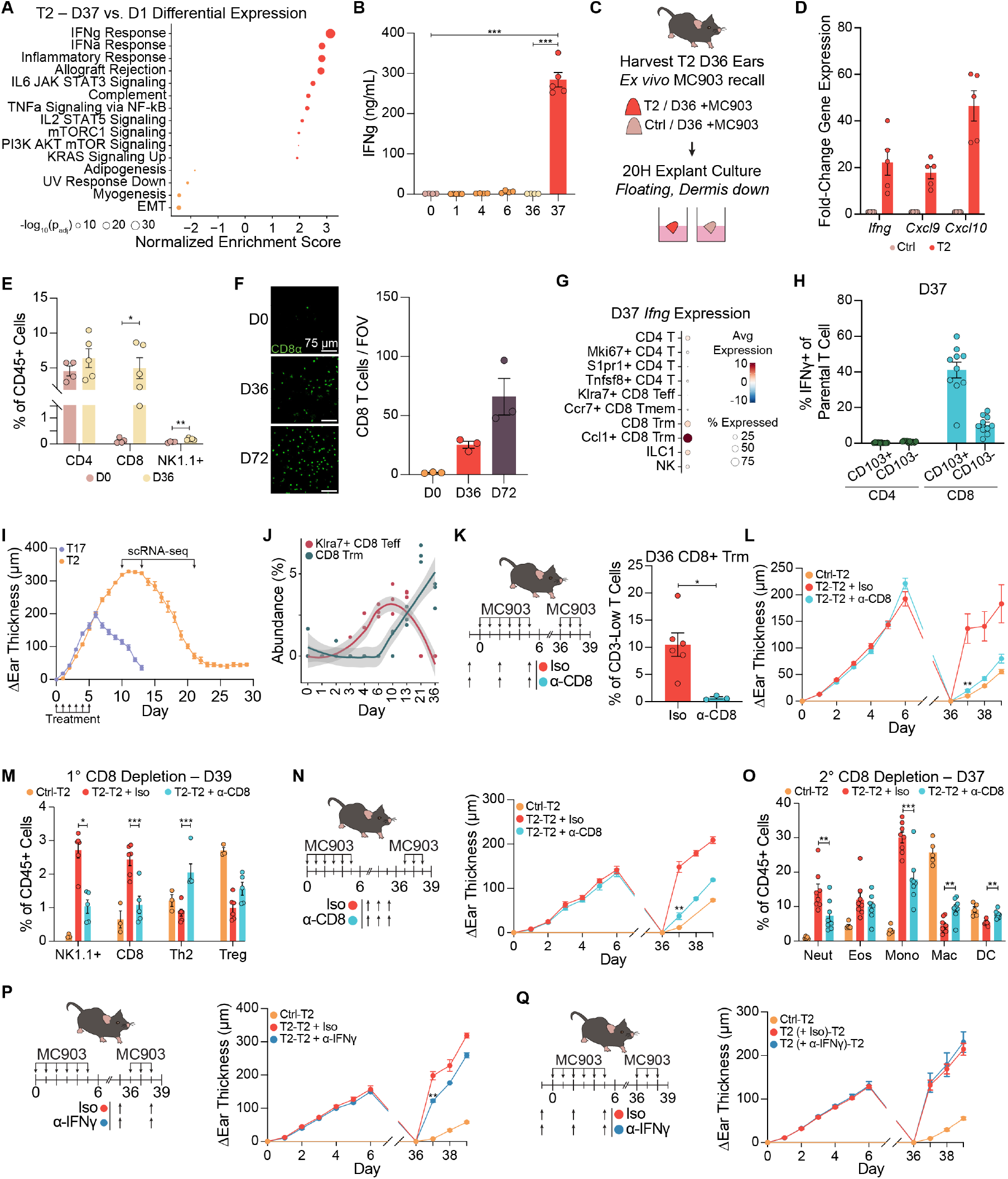
CD8+ Trm-derived IFNγ separates memory recall from acute T2 inflammation. Gene set enrichment analysis (GSEA) was conducted on differentially expressed genes determined by pseudobulk on mouse replicates using all cells captured on D1 and D37 in T2 inflammation. **(B)** Concentration of IFNγ measured in skin lysates during T2 inflammation (n=5 (D1, D37), 4 (D0, D4, D6, D36) mice). **(C)** Schematic of ear skin explants from T2-experienced (primary MC903) and control (primary vehicle) ears. Mice were euthanized, MC903 was applied ex vivo and allowed to dry before harvesting, splitting and culturing for 20 hours. **(D)** qPCR analysis of *Ifng* and ISGs from ear skin explants normalized to control ears (n=5 explants/condition). **(E)** Flow cytometric analysis of abundance of CD4+, CD8+ and NK1.1+ cells in D36 recovered skin (n=4 (D0), 5 (D36) mice). **(F)** Epidermal whole mounts of primary D36 and secondary D72 T2-recovered skin. 5 fields of view (FOV) were quantified across 3 mice/timepoint. Dot plot of *Ifng* expression scaled by z-score in D37 skin. Flow cytometric analysis of IFNγ+ T cells in D37 skin (n=10 mice). **(I)** Longitudinal ear thickness curve tracking recovery after end of 6-day treatment window (n=5 mice). Additional scRNA-seq timepoints for T2 inflammation are denoted. **(J)** Abundance curves for *Klra7*+ CD8 Teff and CD8+ Trm cells across primary T2 inflammation with additional recovery phase timepoints D10-21. **(K)** Schematic of CD8+ T cell depletion in primary T2 inflammation. CD103+ CD8+ Trm cells were analyzed in D36 recovered skin (n=6 (Iso), 3 (α-CD8) mice). **(L)** Ear thickness curve in allergic memory recall (n=5 (Ctrl-T2, α-CD8), 4 (Iso) mice) and **(M)** immune cell representation on D39 after CD8+ T cell depletion in T2 primary challenge (n=3 (Ctrl-T2), 6 (Iso), 5 (α-CD8) mice). **(N)** Schematic of circulating CD8+ T cell depletion before memory recall with ear thickness curve (n=5 (Ctrl-T2), 6 (Iso, α-CD8) mice). **(O)** Immune cell populations in D37 skin after circulating CD8+ T cell depletion (n=5 (Ctrl-T2), 6 (Iso, α-CD8) mice). **(P-Q)** Schematic and ear thickness curves of IFNγ neutralization in **(P)** secondary (n=5 (Ctrl-T2, Iso), 6 (α-IFNγ) mice) and **(Q)** primary T2 challenge (n=5 (Ctrl-T2, α-IFNγ), 10 (Iso) mice).

Given the site-specificity of allergic memory (Fig 1E), we next asked whether IFNγ originated from a tissue-resident source. To eliminate contributions from circulating cells, D36 recovered mice were euthanized and ears were treated with MC903 immediately prior to harvesting. After a 1-minute period for MC903 to dry, ears were collected, split and cultured as skin explants for 20 hours (Fig 3C). Comparing the T2 response between D36 recovered and vehicle-treated explants revealed robust and specific induction of *Ifng* and IFN-stimulated genes (ISGs) like *Cxcl9* and *Cxcl10* in T2experienced ears (Fig 3D). A skin-resident source therefore initiates an IFNγ response that differentiates allergic recall from primary T2 responses.

Next, we hypothesized that the source of IFNγ would be expanded in D36 recovered skin relative to naive skin. Flow cytometric profiling of known IFNγ-producing populations revealed that CD8+ T cells and NK1.1+ cells (comprising ILC1s and NK cells) were expanded on D36 (Fig 3E). Relative to a modest 2.5-fold increase in NK1.1+ cells, CD8+ T cells expanded nearly 40-fold between D0 and D36. Flow cytometry revealed that CD8+ T cells in D36 skin were predominantly CD103+ CD69+ Trm cells (Extended Data Fig 5B). Whole-mount imaging of the epidermis–the principal niche of skin CD8+ Trm cells^48^–clearly demonstrated the presence of Trm cells on D36 (Fig 3F). Interestingly, Trm cells further expanded by approximately 2.5-fold after recovery from secondary challenge (Fig 3F), coinciding with progressively exacerbated one-day recall responses in tertiary T2 challenge (Extended Data Fig 5C).

Finally, we asked whether these CD8+ Trm cells produced IFNγ in recall. On D37, scRNA-seq revealed that *Ccl1*+ CD8+ Trm cells were the dominant source of *Ifng* (Fig 3G). This population represents an activated Trm state that emerged specifically during recall alongside a proportional decline in the resting CD8+ Trm state (Extended Data Fig 5D). By flow cytometry, 40% of CD103+ CD8+ Trm cells produced IFNγ compared to 10% of CD103-CD8+ T cells and <1% of CD4+ T cells (Fig 3H). The geometric mean fluorescence intensity (gMFI) of IFNγ was also substantially higher in CD8+ Trm cells than in other CD4+, CD8+ or NK1.1+ populations (Extended Data Fig 5E). Altogether, these findings position CD8+ Trm cells as the dominant source of IFNγ during T2 memory recall.

To examine the kinetics of Trm cell establishment following T2 primary challenge, we tracked ear thickness throughout the 30-day recovery period. We observed that T2 inflammation self-sustained for 5-6 days after the final MC903 treatment (D5) before gradually resolving (Fig 3I). By contrast, T17 inflammation resolved rapidly following IMQ cessation (Fig 3I). While continuous MC903 treatment is known to sustain ear thickening over a two-week period ^49^, this self-amplifying phase after treatment cessation is not well described. Histologically, this response featured epidermal hyperplasia and immune infiltration rather than edema (Extended Data Fig 5F). Hypothesizing that the prolonged inflammatory phase captured the establishment of CD8+ Trm cells, we further profiled peak inflammation and resolution (D10, D13 and D21) by scRNA-seq (Fig 3I). While *Klra7*+ CD8+ Teff cell abundance closely mirrored ear thickness from D0D21, CD8+ Trm cells first emerged at D10 and progressively expanded during resolution (Fig 3J). The late self-sustaining phase of primary T2 inflammation thus supports seeding of CD8+ Trm cells.

Next, we tested whether CD8+ Trm cells functionally contributed to the T1 inflammatory shift in allergic memory. As epidermal CD8+ Trm cells are not efficiently depleted by peripherally administered antibodies ^50^, we blocked Trm establishment by depleting circulating CD8+ T cells in primary challenge (Fig 3K). CD8+ T cell depletion had no effect on ear swelling in primary T2 inflammation, suggesting that CD8+ T cells are dispensable during the acute response (Extended Data Fig 5G). By contrast, loss of CD8+ Trm cells reduced one-day ear thickening by 85% during secondary challenge (Fig 3K-L). Trm deficiency also reduced infiltration of canonical T1 features like neutrophils, NK1.1+ cells and CD8+ T cells on D39 (Fig 3M, Extended Data Fig 5H). Representation of T2-associated effectors like eosinophils and Th2 cells was concurrently increased (Fig 3M, Extended Data Fig 5H). Altogether, CD8+ Trm cells initiate a T2-to-T1 shift that drives exacerbated ear swelling as part of a tissue response rewired by the encoding of allergic memory.

CD8+ Trm cell deficiency also reduced recruitment of circulating CD8+ T cells during recall, a relationship first nominated by our T2 co-occurrence network (Fig 3M, 2J). To ask how these recruited cells contributed to recall, we depleted CD8+ T cells immediately prior to secondary challenge. This approach eliminated the circulating CD8+ T cell pool without affecting skin CD8+ Trm cells (Fig 3N, Extended Data Fig 5I). Loss of circulating CD8+ T cells produced a 75% reduction in one-day ear thickening on D37 (Fig 3N). Although substantial, the lesser magnitude of this reduction relative to the absence of CD8+ Trm cells suggests a secondary contribution to recall (Fig 3L). While eosinophil abundance was unaffected in the absence of recruited CD8+ T cells, infiltrating neutrophils and monocytes were greatly reduced (Fig 3O). These findings thus support a hierarchical model in which CD8+ Trm cells initiate a T1 shift in T2 allergic memory recall, while infiltrating CD8+ T cells amplify a downstream T1 response.

Finally, we asked whether Trm-derived IFNγ contributed functionally to T2 recall. Antibody-based neutralization of IFNγ immediately before rechallenge reduced one-day ear thickening by 40% (Fig 3P). By contrast, IFNγ neutralization during primary challenge did not alter ear thickening or memory recall after recovery (Fig 3Q, Extended Data Fig 5J). Collectively, our studies establish CD8+ Trm cell-derived IFNγ as a critical initiator of the T2-to-T1 shift that transforms allergic recall from a predominantly T2 response into a mixed T1/T2 response.

### Mouse and human allergic CD8+ Trm cells produce T1 and T2 effectors

To characterize allergic CD8+ Trm cell states in memory, we sorted CD8+ T cells after T2 recovery (D36) and one day of recall (D37). We performed 5’ scRNA-seq with Trm cell surface marker profiling on 10 mice per timepoint (Fig 4A). We captured 7,708 cells spanning two major classes of CD8+ T cells: (1) skin-resident Trm cells, composed almost entirely of *Tbx21*+ T1 CD8+ Trm (Trm1) cells with a small fraction of *Rorc*+ Trm17 cells^51^, and (2) recruited effector (Teff) and memory (Tmem) subsets that emerged during recall (Fig 4B, Extended Data Fig 6A, Supplementary Table 2). We identified 12 transcriptionally distinct states that were well represented across mice within each timepoint (Extended Data Fig 6B-C). CITE-seq confirmed the expected expression of residency markers on Trm subsets (Extended Data Fig 6D).

**Figure 4.**
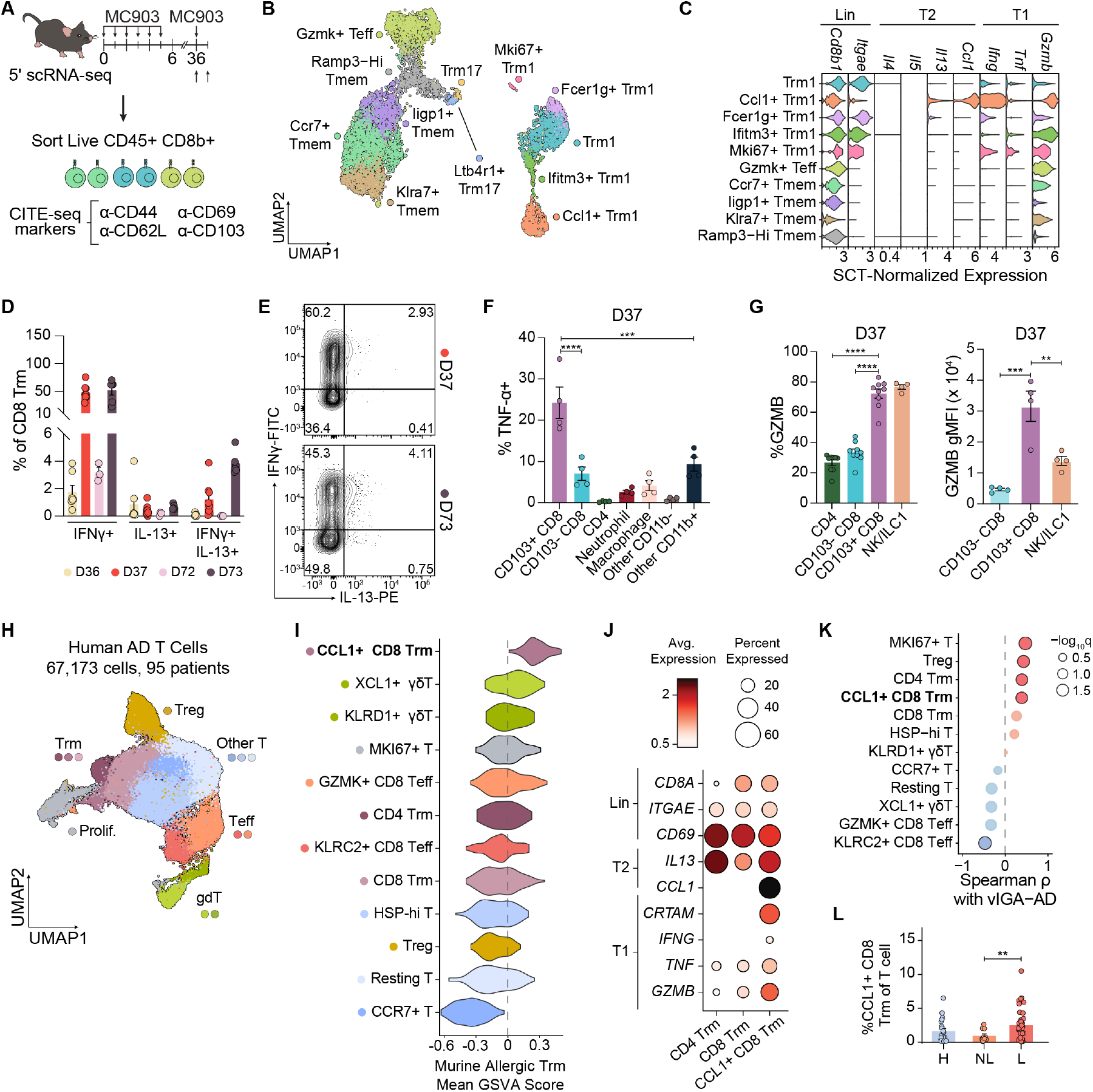
Allergic CD8+ Trm cells produce T1 and T2 effector cytokines that resemble T cell subsets in human AD. **(A)** Schematic of 5′ scRNA-seq of sorted CD8+ T cells from D36 and D37 skin to capture Trm, Teff and Tmem cells. **(B)** UMAP embedding of 7,708 CD8+ T cells across 20 mice (n = 10/timepoint) coloured by cell state. **(C)** Violin plot depicting expression levels of CD8+ T cell lineage markers along with T2 and T1 effector molecules across all CD8+ T cell states. **(D)** Intracellular staining of IFNγ and IL-13 across CD103+ CD8+ Trm cells on D36, D37, D72 and D73 skin (n=7 (D36), 8 (D37), 3 (D72), 6 (D73) mice) with **(E)** representative flow plots for D37 and D73. **(F)** Flow cytometric quantification of TNF-α production across immune cell types on D37 (n=4 mice). **(G)**Quantification of abundance and gMFI of *GZMB* expression across lymphoid populations in D37 (n=9 (% *GZMB*), 4 (gMFI) mice). **(H)** UMAP embedding of 67,173 T cells across 95 patients and 5 studies integrated using Harmony where cell subtypes are annotated and cell clusters are coloured. Study information and clustering details are provided in Extended Data Fig 6A-C. **(I)**Mapping of top 100 genes defining murine allergic *Ccl1*+ CD8+ Trm state as a gene module across human T cell clusters using patient-level pseudobulk GSVA. **(J)** Dot plot of lineage-defining, T2 and T1 effector molecule expression across Trm cell subsets. **(K)** Cell cluster abundances were correlated against Validated Investigator Global Assessment for Atopic Dermatitis (vIGA-AD) scores. Significant associations are highlighted. **(L)** Quantification of *CCL1*+ CD8+ Trm cells in biopsies across healthy (H), non-lesional (NL) and lesional (L) skin (n=34 (H), 20 (NL), 41 (L)patients).

Given that CD8+ Trm cells initiated the T1 shift during allergic memory recall (Fig 3M), we specifically examined expression of T1 and T2 effectors in Trm1 cells. Most Trm1 states expressed T1-associated *Ifng, Tnf* and *Gzmb*. Unexpectedly, two Trm1 populations–*Fcer1g*+ Trm1 and *Ccl1*+ Trm1–also co-expressed *Il13*, but not *Il4* or *Il5*, alongside these T1 molecules (Fig 4C). While *Fcer1g*+ Trm1 represented 10-30% of Trm1 in recovered skin, re-activated *Ccl1*+ Trm1 represented 50-80% of Trm1 cells on D37. This recall-associated state also expressed higher levels of *Ifng, Gzmb* and *Il13* than *Fcer1g*+ Trm1 (Fig 4C, Extended Data Fig 6C). Activated CD8+ Trm cells thus acquire a mixed T1/T2 effector program during T2 memory recall.

Next, we used flow cytometry to determine whether these mixed effector states were recapitulated at the protein level. First, we asked whether CD8+ Trm cells co-produced IFNγ and IL-13. On D37, 50% of CD8+ Trm cells expressed IFNγ alone, 0.2% made only IL-13 and 1.2% produced both cytokines (Fig 4D-E). As Trm cells expanded with repeated T2 challenge (Fig 3F), we further examined these patterns after one day of tertiary challenge (D73). This co-producing population increased to 4% of CD8+ Trm cells whereas the IFNγ single-positive population was maintained at 50% (Fig 4D-E). These co-producers may thus become increasingly prominent with successive inflammatory episodes. Because scRNA-seq also identified *Tnf* and *Gzmb* as prominent effector molecules expressed by activated Trm1 states (Fig 4C), we next verified TNF-α and *GZMB* production by flow cytometry. CD103+ CD8+ Trm cells were the dominant source of TNF-α in D37 rechallenged skin both by frequency and gMFI (Fig 4F, Extended Data Fig 6E). Similarly, while 70% of both CD8+ Trm and NK1.1+ cells expressed *GZMB* in D37 skin, CD8+ Trm cells exhibited substantially higher *GZMB* gMFI, consistent with greater protein abundance (Fig 4G). Together, these data establish allergic CD8+ Trm cells as a dominant source of T1 effector molecules while revealing their capacity to co-produce T1 and T2 cytokines during recall.

To determine if this re-activated murine allergic CD8+ Trm state was conserved in human disease, we integrated, clustered and re-annotated five published AD scRNA-seq datasets spanning healthy, non-lesional and lesional skin (Extended Data Fig 7A-B) ^46,52–55^. Across 95 patients and 67,173 cells, we identified 12 transcriptionally distinct T cell states that were well represented across individuals (Fig 4H, Extended Data Fig 7C-D). We defined a murine activated *Ccl1*+ CD8+ Trm signature using the top 100 cluster-defining differentially expressed genes. This signature mapped most strongly to a human *CCL1*+ CD8+ Trm subset (Fig 4I). Importantly, *CCL1*+ CD8+ Trm cells were recently identified as a treatment-refractory subset despite long-term dupilumab treatment ^52^. Consistent with our murine data, human *CCL1*+ CD8+ Trm cells expressed not only *IL13* and *CCL1* but also the T1 effector molecules *IFNG, TNF* and *GZMB* (Fig 4J). While prior work highlighted *IL13* and cytotoxicity-associated marker *CRTAM* in this treatment-refractory subset ^52^, its mixed T1/T2 profile–including *TNF* and *IFNG* expression– has not been recognized.

We next asked whether this mixed T1/T2 CD8+ Trm state was associated with AD severity as measured by the Validated Investigator Global Assessment for AD (vIGA-AD) score, a 5-point scale of clinical severity. Correlating T cell state abundances with vIGA-AD values identified *CCL1*+ CD8+ Trm cells among the populations most strongly associated with AD severity (Fig 4K). Furthermore, *CCL1*+ CD8+ Trm cells were significantly enriched in lesional relative to non-lesional skin (Fig 4L, Extended Data Fig 7E). Together, these findings demonstrate that the murine *Ccl1*+ CD8+ Trm population driving the T1 shift in allergic memory is both conserved and clinically relevant in human AD. While AD is classically viewed as a T2-driven disease, chronic lesions exhibit prominent IFNγ-associated signatures whose origins and contributions to pathogenesis have remained unclear ^56,57^. Our findings now identify allergic CD8+ Trm cells as a persistent population that drives a T1 shift in chronic AD.

### Fibroblasts amplify CD8+ T cell responses during allergic recall via IL-15

Next, we sought to identify the cell circuits through which CD8+ Trm cells propagate the T1 shift during allergic recall. Within the T2 inflammatory memory network (Fig 2J), several *Tnc*-Hi iFB states co-occurred with activated *Ccl1*+ CD8+ Trm and recruited *Klra7*+ CD8+ Teff cells during secondary challenge (Fig 5A). *Cxcl5*+ *Tnc*-Hi iFBs were the dominant iFB population in the late stages of primary (D10) and secondary (D39) T2 inflammation (Extended Data Fig 8A). By contrast, *Cxcl9*+ *Tnc*-Hi iFBs emerged specifically in T2 recall (Extended Data Fig 8A). Targeted analysis showed that iFBs expressed receptors for Trm-derived T1 and T2 cytokines including *Ifngr1, Il4ra* and *Tnfrsf1a*, suggesting that they may respond to both inflammatory axes within the T1/T2 recall environment (Fig 5B).

**Figure 5.**
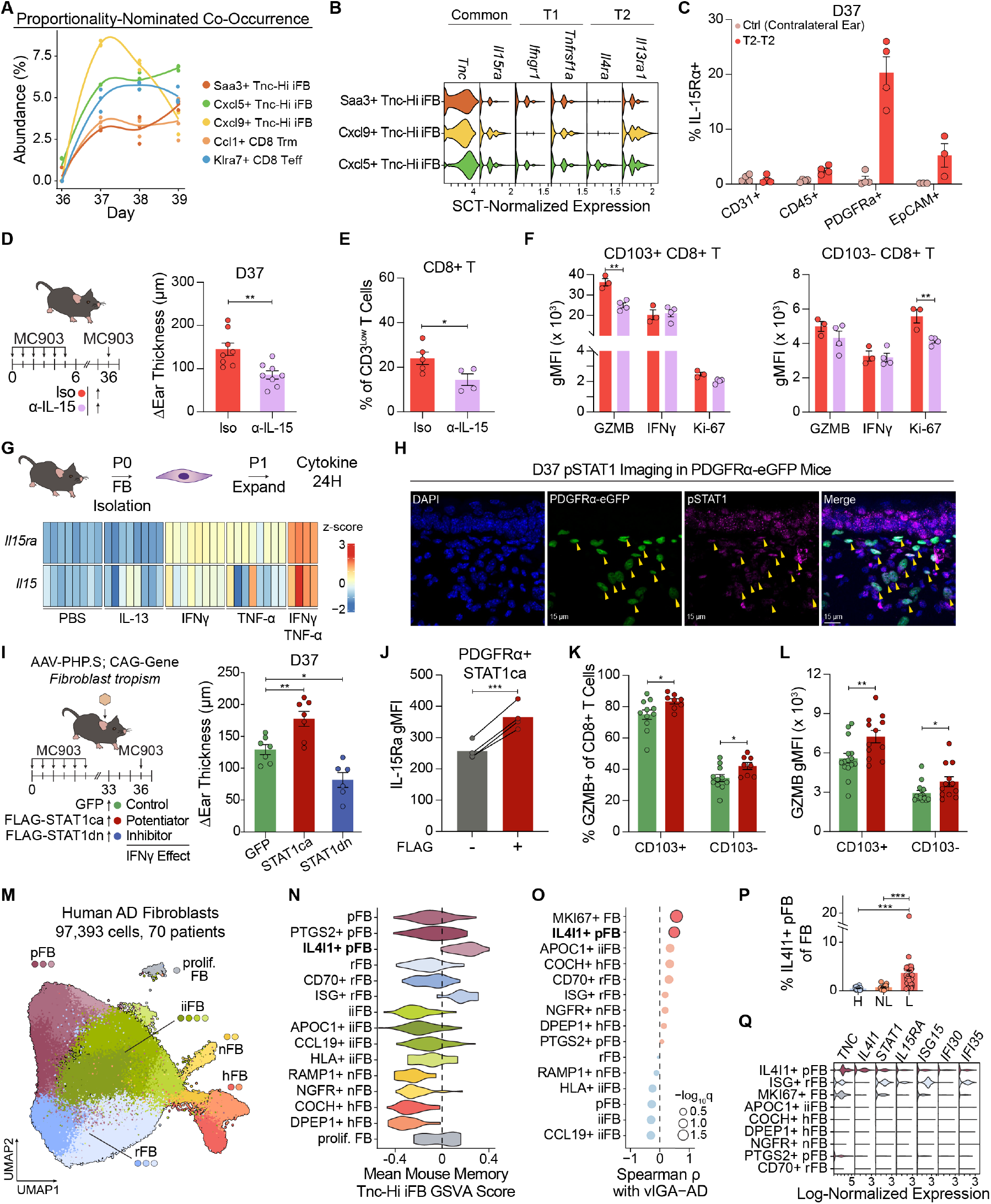
Fibroblasts respond to Trm-derived T1 cytokines to support CD8+ T cells. **(A)** Abundance plots for co-occurring CD8+ T cell and *Tnc*-Hi iFB states nominated by proportionality network in Fig 2J. **(B)** Violin plot showing expression of T1 and T2 cytokine receptors along with selected differentially expressed markers of iFB states. **(C)** Flow cytometry was used to determine IL-15Rα positivity across CD31+ endothelial, CD45+ immune, PDGFRα+ fibroblasts and Ep-CAM+ epithelial cells (n=4 mice). Schematic and D37 ear thickness of IL-15 blockade prior to T2 memory (n=8 mice/group). Abundance of CD8+ T cells in D37 skin measured by flow cytometry (n=5 (Iso), 4 (α-IL-15) mice). **(F)** gMFI of IL-15-mediated support markers *GZMB*, IFNγ and Ki-67 in CD103+ and CD103-CD8+ T cells on D37 (n=3 (Iso), 4 (α-IL-15) mice). **(G)** Schematic of primary ear skin fibroblast isolation for cytokine stimulation. Heatmap of *Il15ra* and *Il15* expression scaled as z-score across cytokine stimulation conditions (n=4 (IFNγ/TNF-α), 8 (single cytokine) mice). **(H)** Representative pSTAT1 staining in D37 skin from PDGFRα-eGFP mice. Yellow arrowheads denote pSTAT1+ PDGFRα+ fibroblasts. Schematic of intradermal AAV administration for *STAT1* signaling modulation prior to T2 recall with associated ear thicknesses on D37 (n=7 (GFP, STAT1ca), 6 (STAT1dn) mice). FLAG-STAT1ca+ PDGFRa+ fibroblasts were compared with FLAG-untransduced fibroblasts for IL-15Rα gMFI on D37 (n=4 mice). **(K)** Frequency of *GZMB* positivity (n=11(GFP), 8 (STAT1ca) mice) and **(L)** *GZMB* gMFI were measured in CD103+ Trm cells and CD103-recruited cells on D37 in AAV-injected mice (n=15 (GFP), 12 (STAT1ca) mice). **(M)** UMAP embedding of 97,393 fibroblasts across 70 patients and 5 studies where subtypes are annotated (reticular rFB, papillary pFB, immune-interacting iiFB, nerve-associated nFB and hair follicle-associated hFB) and clusters are coloured. Study information is provided in Extended Data Fig 6A-B and cluster definitions in Extended Data Fig 9A. **(N)** Mapping of the top 50 genes defining murine *Tnc*-Hi iFB subtype as a gene module across human fibroblast clusters using patient-level pseudobulk GSVA. **(O)** Cell cluster abundances were correlated against vIGA-AD scores. Significant associations are highlighted. **(P)** Quantification of *IL4I1*+ pFB abundance in biopsies across healthy (H), non-lesional (NL) and lesional (L) skin (n=19 (H), 20 (NL), 31 (L) patients). **(Q)** Violin plot of genes stimulated by T2 and T1 effector cytokines across severity-associated human fibroblast clusters.

In addition to *Tnc*, iFB clusters were defined by shared expression of *Il15ra*, the obligate trans-presenting receptor for IL-15 (Fig 5B) ^58^. IL-15 is known to promote CD8+ T cell survival, proliferation and cytotoxicity, but requires transpresentation via IL-15Rα to signal ^58^. Given the *Il15ra* expression, we hypothesized that iFBs reinforce CD8+ T cell responses during allergic recall. At the protein level, fibroblasts robustly upregulated IL-15Rα within one day of secondary challenge (Fig 5C). Approximately 20% of PDGFRα+ fibroblasts expressed IL-15Rα compared to only 5% of epithelial cells and 2.5% of immune cells. Importantly, IL-15Rα induction was minimal in contralateral control ears, indicating that this response is driven by local recall signals rather than systemic factors (Fig 5C).

To determine whether IL-15 trans-presentation functionally contributes to T2 recall, we administered a single dose of IL-15 blocking antibody one day before secondary challenge (Fig 5D). Because IL-15 is required for long-term maintenance of skin Trm cells^59,60^, we employed a short-term blockade strategy that preserved CD8+ Trm cell numbers (Extended Data Fig 8B). Strikingly, IL-15 neutralization reduced one-day ear thickening by roughly 40%, accompanied by a reduction in the proportion of CD8+ T cells (Fig 5D-E, Extended Data Fig 8C). Consistent with its established role in promoting CD8+ T cell effector function ^61^, IL-15 blockade selectively reduced *GZMB* gMFI in CD8+ Trm cells but not in CD103-CD8+ T cells (Fig 5F). The proportion of *GZMB*+ CD8+ Trm cells remained unchanged (Extended Data Fig 8D). We observed no changes in IFNγ expression, but the proliferation marker Ki-67 was selectively reduced in recruited CD8+ T cells (Fig 5F). Together, these findings indicate that iFBs amplify allergic recall by reinforcing IL-15dependent effector responses in both resident and recruited CD8+ T cells. Given that IL-15 is elevated in the dermis of AD lesions ^62^, these findings implicate fibroblast-mediated IL-15 trans-presentation as a clinically relevant mechanism in allergic tissue memory.

Next, we asked whether CD8+ Trm-derived cytokines prime fibroblasts for IL-15 trans-presentation. Primary ear skin fibroblasts were stimulated with IFNγ, TNF-α and IL-13 and profiled by bulk RNA-seq (Fig 5G). *Il15* and *Il15ra* were induced more strongly by IFNγ than TNF-α, and the combination of T1 effector cytokines produced a marked synergistic effect (Fig 5G). Thus, T1 cytokines produced by CD8+ Trm cells establish a feed-forward circuit that amplifies allergic recall through fibroblast-mediated IL-15 trans-presentation. Consistent with this model, imaging of phosphorylated *STAT1* (pSTAT1) in PDGFRα-eGFP reporter mice revealed robust nuclear staining in fibroblasts during recall (Fig 5H). Fibroblasts thus respond to Trm-derived IFNγ in T2 memory.

Because pSTAT1 was also detected in non-fibroblast populations (Fig 5H), we next tested the specific contribution of fibroblast IFN responsiveness to allergic recall. We developed an AAV-based platform to selectively manipulate *STAT1* activity in fibroblasts. Intradermal delivery of 1.5 × 10^11^ gc of an mCherry-encoding AAV transduced approximately 30% of fibroblasts with minimal transduction of other cell types (Extended Data Fig 8E). We generated variants of *STAT1* that were constitutively active (STAT1ca) and dominant-negative (STAT1dn) ^63,64^. To validate these constructs, we generated stable 3T3 fibroblast lines that expressed each *STAT1* variant under doxycycline-inducible control and conducted bulk RNA-seq after IFNγ stimulation (Extended Data Fig 8F). Across IFNγ-responsive genes, STAT1dn inhibited ISG expression by roughly 30% while STAT1ca enhanced responses by 10-60% (Extended Data Fig 8F). Importantly, STAT1ca alone produced minimal transcriptional changes in the absence of IFNγ (Extended Data Fig 8F-G). Thus, STAT1ca potentiates IFNγ responses rather than inducing ISG expression de novo. For in vivo studies, FLAG-tagged *STAT1* variants were placed under a constitutive CAG promoter to enable identification of transduced cells.

To manipulate fibroblast IFN responsiveness in situ, we allowed mice to recover from primary T2 inflammation and intradermally delivered AAV-*STAT1* variants prior to secondary challenge (Fig 5I). After allowing three days for transgene expression, we induced allergic recall (Fig 5I). Potentiation of fibroblast IFN responsiveness with STAT1ca increased one-day ear thickening by approximately 40%, whereas inhibition through STAT1dn produced a reduction of similar magnitude (Fig 5I). We next compared IL-15Rα expression between FLAG+ fibroblasts transduced with STAT1ca and untransduced FLAGneighbours in the same recall environment. Consistent with a potentiation effect, STAT1ca-transduced FLAG+ fibroblasts exhibited significantly higher IL-15Rα gMFI than FLAG-fibroblasts (Fig 5J). Correspondingly, CD8+ T cells in STAT1ca-transduced mice showed increases in both the frequency of *GZMB*+ cells and *GZMB* gMFI across resident and recruited populations (Fig 5K-L). Together, these findings demonstrate that Trm-derived T1 cytokines promote IL-15 trans-presentation by fibroblasts, thereby amplifying cytotoxic CD8+ T cell responses. This bidirectional stromal-immune circuit grounded in a feed-forward motif links Trm re-activation to downstream tissue inflammation in allergic memory recall.

Having identified iFBs as downstream amplifiers of the T1 shift during allergic recall, we finally asked whether analogous fibroblast programs exist in human AD. Using our integrated human dataset, we captured 97,393 fibroblasts across 70 patients and 5 major subtypes: papillary (pFB), reticular (rFB), immune-interacting (iiFB), nerve-associated (nFB) and hair follicle-associated (hFB) (Fig 5M). Iterative clustering identified 15 distinct states that were well represented across patients (Fig 5M, Extended Data Fig 9A-B). We generated a murine iFB signature using the top 50 differentially expressed genes between D37 and D36 fibroblasts to capture the recall-associated iFB program (Extended Data Fig 8A). Mapping this signature onto the human atlas revealed the strongest enrichment within *IL4I1*+ pFBs and ISG+ rFBs (Fig 5N). Of these two populations, only *IL4I1*+ pFBs were significantly associated with vIGA-AD-scored clinical severity (Fig 5O). Finally, this state was enriched in lesional compared with non-lesional skin (Fig 5P, Extended Data Fig 9C).

Previous studies have described this AD-associated *IL4I1*+ pFB population as a T2-responsive fibroblast state based on expression of *IL4I1* and *TNC* ^46,55,65^. However, GSEA of state-defining genes also revealed significant enrichment of IFNγ and *TNF* signaling pathways within *IL4I1*+ pFBs (Extended Data Fig 9D). Notably, *IL4I1*+ pFBs expressed not only known markers of T2 responsiveness (*IL4I1, TNC*) but also T1 responsiveness (*STAT1, ISG15*) (Fig 5Q). Furthermore, these cells expressed *IL15RA*, revealing a conserved capacity for IL-15 trans-presentation that aligns with elevated IL-15 levels in the dermis of AD lesions (Fig 5Q) ^62^. Taken together, we identify a conserved IFN-responsive fibroblast program in human AD that mirrors iFBs in allergic memory. The fibroblast-CD8+ T cell circuit that amplifies allergic recall responses may thus be conserved in AD patients.

### Fibroblast-derived chemokines promote CD8+ T cell recruitment

iFBs co-occurred with *Ccl1*+ CD8+ Trm cells and recruited CD8+ Teff cells during allergic recall (Fig 5A). Given our hierarchical model in which CD8+ Trm cells initiate recall and recruited CD8+ T cells amplify the response, we hypothe-sized that iFBs bridge these two phases through CD8+ T cell recruitment. CellChat analysis of iFB-CD8+ T cell interactions during T2 memory nominated two candidate chemokine axes: *Cxcl12*-Cxcr4 and *Ccl19*-*Ccr7* (Fig 6A). While *Cxcl12* was expressed by both iFBs and endothelial cells, *Ccl19* was produced exclusively by iFBs (Fig 6B). Protein measurements in skin lysates revealed distinct kinetics, with CXCL12 predominating in primary T2 inflammation whereas *CCL19* was uniquely induced during recall (Fig 6C). In the human atlas, *CCL19* expression was also elevated in lesional AD fibroblasts (Fig 6D). These results suggest that CXCL12 supports CD8+ T cell recruitment during primary T2 inflammation, while *CCL19* supports CD8+ T cell recruitment during recall.

**Figure 6.**
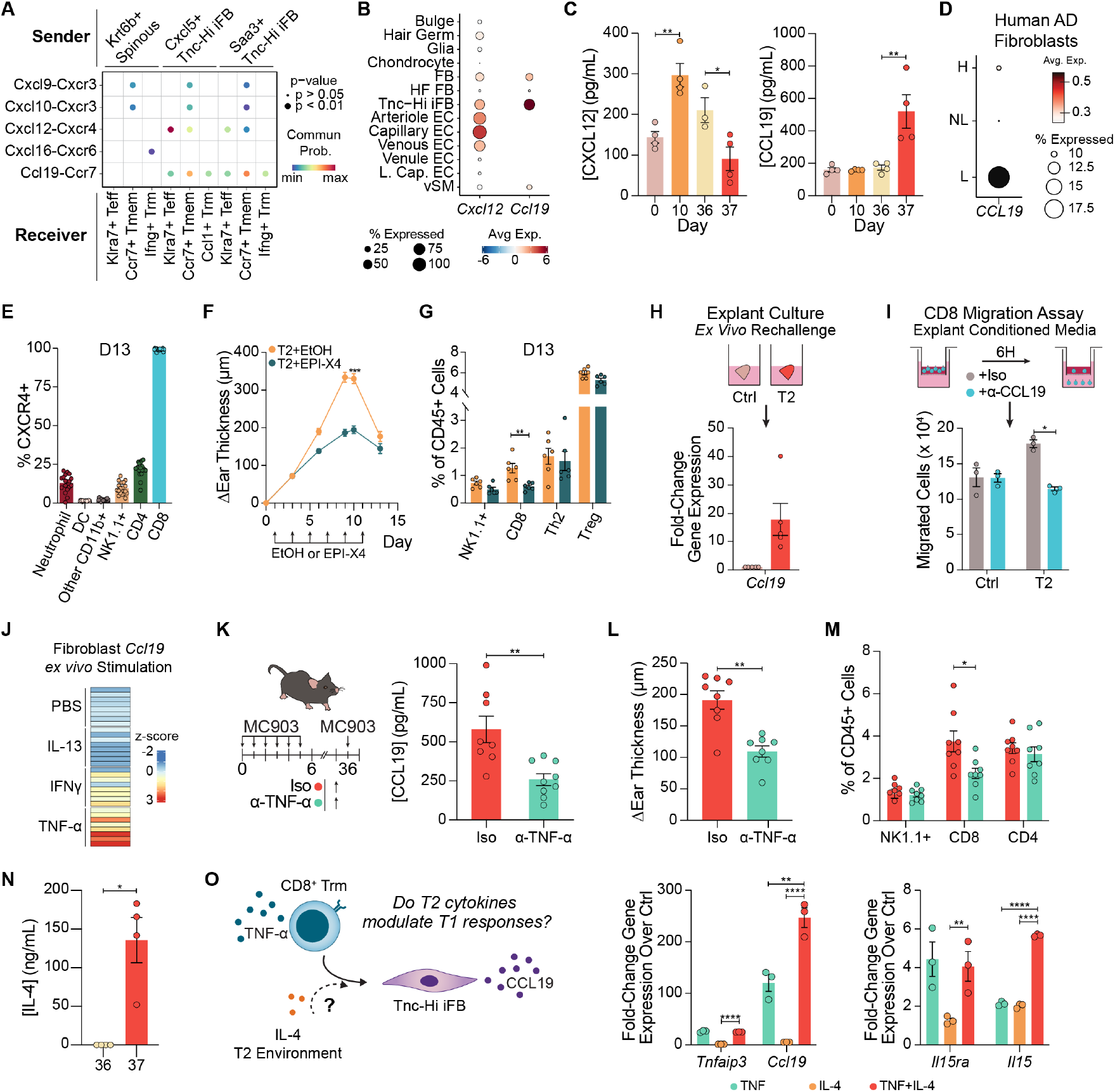
Fibroblasts responding to CD8+ Trm-derived TNF-α recruit CD8+ T cells via *CCL19*. **(A)**CellChat cell-cell communication analysis of chemokine pathways between structural cells and CD8+ T cell states nominated by proportionality analysis in Fig 2J. **(B)** Dot plot of *Cxcl12* and *Ccl19* expression scaled by z-score in T2 single-cell timecourse. **(C)** CXCL12 and *CCL19* protein levels in ear skin lysates across primary and secondary MC903. **(D)** Dot plot of single-cell fibroblast *CCL19* expression across healthy (H), AD non-lesional (NL) and AD lesional (L) human skin. **(E)** Proportion of CXCR4+ cells across immune cell subtypes present in skin on D13 of T2 inflammation (n=18 mice). **(F)** Ear thickness curve of primary MC903 with treatment by EPI-X4 to block CXCR4-mediated recruitment (n=25 mice/condition). EPI-X4 or ethanol vehicle-control treatment days are denoted by arrows. **(G)** Immune cell representation in D13 skin after EPI-X4 treatment (n=6 mice/condition). **(H)** Explant schematic and *Ccl19* qPCR after 20-hour culture (n=5 explants). **(I)** Schematic of Transwell CD8+ T cell migration assays to explant-conditioned media with antibody-mediated *CCL19* blockade. Migrated CD8+ T cells in the lower chamber were counted by flow cytometry (n=3 explants/condition). **(J)** Heatmap of *Ccl19* expression by primary fibroblasts after Trm-derived cytokine stimulation measured by bulk RNA-seq (n=8 mice/condition). **(K)** Schematic of TNF-α blockade prior to T2 recall with ELISA measurement of *CCL19* levels on D37 in ear skin lysates (n=8 (Iso), 9 (α-TNF-α) mice). **(L)** One-day ear thickness changes with TNF-α blockade (n=8 mice/condition). **(M)** Quantification of abundance of NK1.1+ cells and T cells in D37 skin after TNF-α blockade (n=8 mice/condition). **(N)** Concentration of IL-4 in skin lysates in T2 memory (n=4 mice/timepoint). **(O)** Left: Schematic illustrating hypothesized interaction between IL-4 and TNF-α signaling in iFBs. Right: qPCR analysis of primary dermal fibroblasts stimulated with IL-4, TNF-α or the combination for 48 hours (n=3 mice/condition).

First, we investigated whether the CXCL12-CXCR4 axis supports CD8+ T cell recruitment in primary T2 inflammation. Nearly all skin-infiltrating CD8+ T cells expressed CXCR4 on D13 (Fig 6E). Functionally, topical administration of the CXCR4 antagonist EPI-X4^66^ throughout primary T2 challenge not only suppressed peak D10 ear swelling by 45%, but also markedly reduced CD8+ T cell infiltration at D13, a timepoint corresponding to the onset of CD8+ Trm formation (Fig 6F-G, Fig 3J). By contrast, the abundance of NK1.1+ and Th2 cells remained unchanged (Fig 6G). These findings identify stromal CXCL12 as a key mediator of CD8+ T cell recruitment during the primary inflammatory phase that coincides with CD8+ Trm establishment (Fig 3J).

Next, we turned to the memory-specific *CCL19*-CCR7 axis. Using our explant system, we observed robust *Ccl19* induction in explants from T2-experienced ears at D36, mirroring *CCL19* protein induction on D37 (Fig 6H, 6C). We next tested whether conditioned media from these explants, which model primary (control) or recall (T2-experienced) responses ex vivo, promoted CD8+ T cell chemotaxis in transwell migration assays (Fig 6I). Conditioned media from T2-experienced ears promoted a 1.5-fold increase in CD8+ T cell migration relative to conditioned media from contralateral control ears (Fig 6I). Strikingly, *CCL19* neutralization abolished this enhanced migration, restoring chemotaxis to levels observed for contralateral controls (Fig 6I). These findings indicate that fibroblast-derived *CCL19* promotes CD8+ T cell migration specifically in the setting of allergic recall.

Having established *CCL19* as a recall-specific chemokine, we next asked whether CD8+ Trm activation induced its production by fibroblasts. Among the Trm-derived cytokines, TNF-α was identified as a potent inducer of *Ccl19* in ex vivo cultured fibroblasts by bulk RNA-seq (Fig 6J). We therefore tested whether TNF-α drives *CCL19* induction in vivo. TNFα neutralization prior to secondary challenge markedly reduced *CCL19* protein levels in D37 skin (Fig 6K). Concurrently, we observed a 40% reduction in one-day ear thickening and diminished CD8+ T cell recruitment (Fig 6L-M). No changes were observed in the abundance of CD4+ T cells or NK1.1+ cells (Fig 6M). Together, these findings identify TNF-α-induced *CCL19* production as a fibroblast-mediated mechanism that recruits CD8+ T cells. This feed-forward motif between fibroblasts and CD8+ T cells thus reinforces the T1 shift in allergic recall.

Although we initially focused on how fibroblasts respond to T1 cytokines during allergic recall, fibroblast IL-4 responses are known to support acute T2 inflammation ^27,45^. Our memory iFBs expressed *Il4ra* and were marked by *Tnc*, a known IL-4 response gene (Fig 5B) ^65^. Importantly, IL-4 was strongly induced on D37, consistent with the T2 exaggeration of recall (Fig 6N, Fig 2H-I). We therefore asked whether IL-4 in the mixed T1/T2 recall environment modulated fibroblast responses to T1 cytokines (Fig 6O). In ex vivo cultured ear skin fibroblasts, IL-4 greatly potentiated *Ccl19* induction by TNF-α despite minimal induction by IL-4 alone (Fig 6O). This synergy was selective, as IL-4 did not affect the canonical TNF-α response gene *Tnfaip3* (Fig 6O). Because Trmderived T1 cytokines also drove *Il15* and *Il15ra* expression, we next asked whether IL-4 similarly potentiated the IL-15 arm of the fibroblast-CD8+ T cell circuit. IL-4 enhanced TNF-α-mediated *Il15* induction but had no effect on *Il15ra* (Fig 6O).

These findings suggest that the mixed T1/T2 signatures observed in murine iFBs (Fig 5B) and human *IL4I1*+ pFBs (Fig 5Q, Extended Data Fig 9D) likely reflect functional synergy between T1 and T2 cytokine programs in vivo. The integrated response to Trm-derived T1 signals and the IL-4-rich tissue milieu thus enhances the capacity of dermal fibroblasts to support allergic memory recall across the IL-15 and *CCL19* axes.

## Discussion

To define the organizational principles of memory across Types of inflammation, we adapted T17 and T2 murine models of skin inflammation into repeated challenge paradigms. Consistent with the prevailing model of Type-specific exaggeration, T17 memory was characterized by accelerated reemergence of the multicellular network established during primary challenge. By contrast, allergic T2 memory departed from this organizational principle and elicited a more severe and durable recall response in which exaggerated T2 inflammation was layered with prominent T1 immune features. This T2-to-T1 shift was initiated by allergic CD8+ Trm cells that produced both T1 and T2 cytokines and mapped onto a treatment-refractory Trm subset in AD skin. Responding to this mixed inflammatory environment, fibroblasts amplified allergic recall through reciprocal interactions with CD8+ T cells involving IL-15 trans-presentation and *CCL19*-mediated recruitment (Extended Data Fig 10). Our findings demonstrate that allergic inflammatory memory departs from the prevailing paradigm of Type-specific exaggeration by rewiring tissue circuitry to promote a mixed T1/T2 ecosystem in recall.

Through unbiased longitudinal profiling of multicellular tissue networks, our compositional scRNA-seq framework revealed distinct organizational principles underlying memory responses. T17 responses were organized around epithelialimmune and epithelial-stromal interactions involving IL-17responsive keratinocytes, γδT cells, neutrophils and fibroblasts. Consistent with Type-specific exaggeration, T17 recall exhibited increased *Il17a*+ γδT cell abundance along-side accelerated re-emergence of the same multicellular network established during primary challenge. These findings demonstrate that Type-specific exaggeration in T17 memory is not simply the product of expanded pathogenic lymphocytes ^16^, but rather a coordinated multicellular response that aligns with proposed models of psoriasis pathogenesis ^36,41^. Conversely, allergic memory generated a larger and more interconnected network that united canonical T2 effectors (basophils), newly recruited T1 populations (pDCs, NK cells and CD8+ T cells) and structural cells (fibroblasts). Rather than multicellular networks of this scale, many T2 acute ^12,13,31^ and memory ^14^ responses have previously been grounded in discrete two-cell circuits supported by positive feedback motifs ^9^. Allergic memory in AD thus reprograms the tissue ecosystem to introduce a new T1 axis whereas T17 memory reinforces the primary inflammatory network.

CD8+ Trm cells capable of producing T1 and T2 effectors like IFNγ, TNF-α and IL-13 emerged as a central node in allergic memory. Re-activation of these allergic Trm cells drove exacerbated one-day ear swelling in secondary challenge. However, both CD8+ T cells and IFNγ signalling were dispensable in primary T2 inflammation. The recruitment and functional contributions of T1 immune features are thus a property of allergic memory rather than an intrinsic feature of T2 responses. Critically, loss of allergic CD8+ Trm cells drastically altered T2 memory by inhibiting the T2-to-T1 shift in recall, restoring a response that instead resembled canonical Type-specific exaggeration. Under these conditions, recruitment of T1 effectors was drastically reduced, while Th2 cell and eosinophil abundances increased. These findings establish allergic CD8+ Trm cells as a principal reservoir of inflammatory memory whose re-activation rewires the tissue ecosystem during secondary challenge.

Our findings also provide a mechanistic explanation for longstanding observations of T1 immunity in AD. Elevated IFNγ and TNF-α signatures have been described in chronic lesional skin, yet their cellular origins and functional roles have remained unclear ^56,57^. Although a treatment-refractory *IL13*+ *CRTAM*+ CD8+ Trm population was recently identified in resolved AD skin ^52^, it was not recognized as a source of T1 cytokines. Using our integrated human AD atlas, we re-identified these cells as *CCL1*+ CD8+ Trm, established their correlation with increasing disease severity, and demonstrated strong alignment with murine allergic Trm cells. Across both species, allergic CD8+ Trm cells expressed IFNγ, TNF-α and IL-13–a previously unrecognized mixed T1/T2 phenotype. While prior work emphasized the capacity of IFNγ to suppress acute T2 responses^67,68^, our findings instead demonstrate that T1 cytokines can amplify allergic memory. Beyond AD, increased IFNγ signaling has also been reported in allergic contexts like severe asthma and eosinophilic esophagitis ^69,70^. Progressive acquisition of T1 immunity may thus represent a common mechanism that sustains pathologic inflammatory memory across chronic allergic disease. Defining when and how T1 and T2 programs interact over recurrent inflammatory episodes may therefore refine current T2-centric models of allergic disease pathogenesis.

Importantly, our compositional analyses demonstrated that allergic memory is not recalled solely through CD8+ Trm cells, but instead through a broader multicellular tissue network. iFBs emerged as key partners of allergic CD8+ Trm cells, forming a reciprocal amplification circuit during recall. Fibroblasts responded to Trm-derived T1 signals by inducing an IL-15 trans-presentation program that enhanced proliferative and cytotoxic features of CD8+ T cells. Although IL15 is an established homeostatic survival factor for skin Trm cells^59,60^, its elevation in AD lesions suggests an additional role in allergic inflammation. We further found that T1 and T2 effector cytokines synergistically induced *Il15* expression in fibroblasts. Because induction of *Il15* and *Il15ra* can occur independently of IL-4, this arm of the fibroblast-CD8+ T cell circuit may not be effectively targeted by T2-directed treatments in AD. *GZMB* may also contribute to AD pathogenesis by cleaving epithelial barrier proteins, thereby compromising barrier integrity ^71,72^. Therapeutically targeting IL-15 or downstream cytotoxic effectors such as *GZMB* may thus provide a strategy to disrupt the T1 circuitry that emerges during allergic memory recall.

To directly interrogate the contribution of fibroblasts to the allergic T1 shift, we developed an AAV-based platform for in situ genetic manipulation of dermal fibroblasts. Unlike depletion approaches ^73^, this system enabled selective perturbation of fibroblasts while preserving their essential contributions to tissue structure. Using this platform, we modulated *STAT1* function during secondary challenge to study fibroblast-intrinsic IFNγ signalling. Enhancing the *STAT1* response downstream of Trm-derived IFNγ was sufficient to exacerbate allergic memory, whereas inhibiting *STAT1* suppressed recall. We further linked the potentiation of *STAT1* signaling to increased IL-15Rα expression by transduced fibroblasts along with enhanced *GZMB* expression in resident and recruited CD8+ T cells. Together, these findings identify fibroblast-intrinsic T1 signaling as a key mechanism by which the fibroblast-CD8+ T cell circuit amplifies allergic recall. Future work can adapt the genetic cargo ^74^ in our AAV-based platform to study iFB-intrinsic responsiveness and further dissect their organizational properties within skin circuitry.

In allergic recall, we defined a hierarchical model in which CD8+ Trm cells initiate T1 responses that are subsequently amplified by recruited CD8+ T cells. Fibroblasts bridged these two phases of recall by producing *CCL19* in response to Trm-derived TNF-α, thereby promoting CD8+ T cell recruitment into skin. Although *CCL19* is classically associated with immune trafficking in secondary lymphoid organs ^75^, emerging work has expanded its known functions to include CD8+ T cell recruitment into the brain and lung during viral infection ^76,77^. Beyond extending this function to the skin, we found that IL-4 selectively potentiated TNF-α-driven *Ccl19* expression in fibroblasts. Our data thus position fibroblast-derived *CCL19* as a point of convergence between T1 and T2 immunity during allergic memory. This responsiveness may further differentiate the role of fibroblasts in allergic memory from viral infection. Furthermore, human iFBs exhibited elevated *CCL19* expression together with responsiveness to both T1 and T2 cytokines. Increased *CCL19* has also been reported in chronic rhinosinusitis ^78^, suggesting that this axis may extend to other chronic allergic diseases. Whether fibroblasts themselves can intrinsically store allergic inflammatory memory or function primarily as response amplifiers is an important question for future study.

Altogether, our work broadens the conceptual framework of inflammatory memory by demonstrating how distinct inflammation Types engage unique multicellular tissue programs during recall. Re-engagement of encoded memories can range from canonical Type-specific exaggerations grounded in identical cellular circuitry to largely rewired tissue responses that drive the acquisition of new immune axes. Our comparative framework further provides an experimental platform for investigating how chronic inflammatory diseases like psoriasis and AD may reinforce or diverge from the immune responses that drove their initial presentation. We establish that inflammatory memory is a tissue-level property that emerges from coordinated responses between immune and structural cells. Our work adds to emerging studies identifying lymphocyte-structural cell circuits in inflammatory settings ^12,24,31^. Through its ability to reorganize tissue circuitry during memory recall, we propose that this immune-stromal motif may be central to stabilizing chronic inflammatory disease in tissue ecosystems. Future studies of inflammatory memory across tissue ecosystems and immune Types will further define the principles underlying the recurrence and progression of chronic inflammatory disease.

## Acknowledgements

We thank all members of the Ordovas-Montanes lab for discussions and feedback. I.O. was supported by the Predoctoral Individual National Research Service Award (1F31AR086579). P.L. was supported by the Howard Hughes Medical Institute James H. Gilliam Fellowship for Advanced Study. J.O.M. is a New York Stem Cell Foundation – Robertson Investigator. J.O.M was supported by The Pew Charitable Trusts Biomedical Scholars, The Broad Next Generation Award, The Mathers Foundation, The New York Stem Cell Foundation, and The Cell Discovery Network, a collaborative funded by The Manton Foundation and The Warren Alpert Foundation at Boston Children’s Hospital. We thank the Henriques Lab for the bioRxiv template which we used as a base for preparing this preprint.

## Author contributions

A.C.K., I.O. and J.O.M. conceived the study. A.C.K. and I.O. performed most of the experiments. L.Y. assisted with flow cytometry, transwell migration assays and ex vivo fibroblast culture experiments. P.L. generated and validated *STAT1* constructs and assisted with AAV-based experiments. A.C.K., I.O. and J.O.M. wrote the paper. All the authors contributed to data interpretation and manuscript editing.

## Materials and Methods

### Mice

Female C57BL/6J mice (The Jackson Laboratory, Stock 00664) and Pdgfra-eGFP reporter mice (The Jackson Laboratory, Stock 007669) were used between 8 and 12 weeks of age for all experiments unless otherwise indicated. Mice were housed under specific pathogen-free conditions at Boston Children’s Hospital. All animal procedures were approved by the Institutional Animal Care and Use Committee (IACUC) and conducted under the supervision of the Department of Animal Resources at Children’s Hospital (Protocol 00002234).

### Inflammatory memory models

All inflammatory memory models employed a dualchallenge paradigm. Murine ear skin received six consecutive days of primary topical treatment, followed by a 30-day recovery period and a second six-day treatment course. The recovery period was at least 30 days and extended where required by the experimental design. For T2 inflammation, 2 nmol MC903 (Tocris) was applied in 20 µL ethanol (10 µL per side) to the left ear and the contralateral right ear received ethanol vehicle alone ^1^. For T17 inflammation, 25 mg of 5% IMQ cream (Perrigo) was applied to the left ear ^2^. Ear thickness was measured using a digital micrometer (Mitutoyo). Baseline ear thickness was recorded on D0 and D36 before challenge.

Additional variations were performed to characterize the properties of inflammatory memory. Durability was assessed by extending the recovery period to approximately 70 weeks. Site-specificity was evaluated by re-challenging either the previously inflamed left ear (site-matched recall) or the contralateral vehicle-treated right ear (site-unmatched recall). Stimulus specificity was assessed by replacing MC903 with IMQ during secondary challenge. For tertiary MC903 experiments, mice underwent an additional 30-day recovery period after secondary challenge before receiving a third six-day course of MC903.

### Histology and immunohistochemistry

#### Histology

Ear skin was harvested immediately after euthanasia and fixed in 10% neutral-buffered formalin for 24 hours at 4°C. The peripheral margins of the ears were trimmed to retain the central ~ 1 cm of tissue before transferring to 70% ethanol and submitting to the Beth Israel Deaconess Medical Center Histology Core Facility (RRID: SCR_009669) for paraffin embedding, 5-µm sectioning, and staining. Hematoxylin and eosin staining was used to assess epidermal hyperplasia and inflammatory infiltrates. Masson’s trichrome staining was performed to evaluate collagen deposition and fibrotic architecture.

#### Epidermal whole mounts

For epidermal whole-mount immunofluorescence, ears were harvested immediately after euthanasia, and dorsal and ventral halves were separated manually. Ear halves were incubated dermis-side down in pre-warmed 20 mM EDTA in PBS for 45 minutes at 37°C with gentle agitation to facilitate separation of the epidermis from the dermis. Epidermal sheets were carefully peeled from the dermis, fixed in 4% paraformaldehyde for 10 minutes, washed in PBS, and blocked for 1 hour at room temperature in blocking buffer (5% donkey serum, 1% BSA, 2% cold-water fish gelatin and 0.3% Triton X-100 in PBS). Epidermal sheets were then incubated overnight at 4°C with gentle agitation in blocking buffer containing Alexa Fluor 488-conjugated anti-mouse CD8α antibody (1:50 dilution, BioLegend 100726), washed in 0.3% Triton X-100 in PBS, and mounted in ProLong Gold Antifade Mountant (Invitrogen). Whole-mount preparations were imaged within one week of mounting.

#### Immunofluorescence of frozen sections

For immunofluorescence staining of frozen tissue sections, whole ears were fixed overnight in 4% paraformaldehyde at 4°C, washed in PBS, cryoprotected in 30% sucrose at 4°C until tissues sank (1-3 d), and embedded in optimal cutting temperature compound (Sakura Finetek). 40-µm sections were then fixed again in 4% paraformaldehyde for 2 minutes and blocked for 1 hour at room temperature in blocking buffer. Slides were stained with primary antibodies overnight at 4°C. After washing in 0.3% Triton X-100 in PBS, sections were incubated with species-specific secondary antibodies together with DAPI where indicated, mounted in ProLong Gold Antifade Mountant (Invitrogen), and allowed to cure before imaging. For phospho-protein staining, sections underwent methanol permeabilization before blocking.

#### Image acquisition and analysis

Brightfield images were acquired using a Zeiss AxioImager slide-scanning microscope at the Boston Children’s Hospital Cellular Imaging Core (RRID: SCR_026485). Confocal fluorescence images were acquired using a Zeiss LSM 880 laserscanning confocal microscope at the Harvard Digestive Disease Center Cell Function and Imaging Core. Identical image acquisition settings were maintained for all samples within each experiment. Image analysis was performed in ImageJ.

To quantify CD8+ Trm cells in recovered ear skin, CD8α+ cells in epidermal whole mounts were manually counted from four randomly selected fields of view per ear, and counts were averaged to obtain one value per biological replicate.

### Whole-tissue protein quantification

Whole ears were harvested immediately after euthanasia and homogenized in Tissue Extraction Reagent I (ThermoFisher) supplemented with protease inhibitors in Lysing Matrix A tubes (MP Biomedicals). A Bead Ruptor homogenizer was used at 4°C with 4 cycles at 6.0 m/s and a 5-minute rest between cycles. Homogenates were clarified by centrifugation at 14,000 × g for 10 minutes at 4°C, and supernatants were collected and stored at −80°C until analysis. Cytokine quantifications were performed according to the manufacturer’s instructions using the following kits: IFN-α1 ELISA (BioLegend 447904), CXCL12 ELISA (R&D Systems DY460), *CCL19* ELISA (R&D Systems DY440), LEG-ENDplex Mouse Cytokine Panel 2 (BioLegend 741334) and LEGENDplex Mouse T Helper Cytokine Panel 3 (BioLegend 741043).

### Flow cytometry

#### Tissue digestion for flow cytometry

Single-cell suspensions for flow cytometry were prepared from mouse ear skin using digestion protocols optimized for immune, fibroblast or epithelial cell isolation. For immune cell analyses, ears were separated into dorsal and ventral halves and digested in RPMI-1640 medium with 10 mM HEPES, 2 mM GlutaMAX, 0.1 mM non-essential amino acids (NEAA), 1 mM sodium pyruvate, 250 µg/mL Liberase TL (Sigma-Aldrich) and 250 µg/mL DNase I (Roche) for 90 minutes at 37°C and 5% CO2. Tissue was then mechanically dissociated by mincing and triturating prior to filtering through 70-µm cell strainers. For intracellular cytokine staining, brefeldin A was added at 5 µg/mL during tissue digestion and maintained throughout subsequent steps until fixation.

For fibroblast analyses, dorsal and ventral ear halves were minced prior to digestion in DMEM with 10 mM HEPES, 2 mM GlutaMAX, 0.1 mM NEAA, 1 mM sodium pyruvate, 2 mg/mL Collagenase A (Roche) and 100 µg/mL DNase I for 30 minutes at 37°C with agitation. Cells liberated during the first digest were collected by gently scraping the dermis with a scalpel, and the remaining tissue was subjected to a second 30-minute digestion in fresh media before pooling both fractions and filtering through 70-µm cell strainers.

For epithelial analyses, ear halves were incubated epidermis-side down in 0.125% trypsin in HBSS for 30 minutes at 37°C with agitation. Epidermal cells were mechanically separated from the dermis by scraping with a scalpel and filtered through 70-µm cell strainers.

#### Flow cytometry staining and acquisition

Single-cell suspensions were stained with Zombie NIR Fixable Viability Dye (BioLegend 423105) in PBS for 20 minutes at 4°C and subsequently washed with fluorescence-activated cell sorting (FACS) buffer (PBS, 2 mM EDTA and 2% fetal bovine serum (FBS)). Samples were then stained with antibodies against surface antigens diluted in FACS buffer for 20 minutes at 4°C and washed three times with FACS buffer. For intracellular staining, cells were fixed and permeabilized using the eBioscience Foxp3 Transcription Factor Staining Buffer Set (ThermoFisher 00-5523-00) for 30 minutes at 4°C, washed once with permeabilization buffer, and blocked in permeabilization buffer containing 2% rat serum for 10 min. Cells were then incubated overnight at 4°C with antibodies against intracellular antigens diluted in permeabilization buffer, followed by three washes with permeabilization buffer and one wash with FACS buffer. Finally, cells were resuspended in FACS buffer containing 123count eBeads (ThermoFisher 01-1234-42). Flow panels are listed in Supplementary Table 3.

Flow cytometry data were acquired using a BD FACSymphony A5 SE cytometer. Absolute cell numbers were calculated using 123count eBeads according to the manufacturer’s instructions. Data were analyzed using FlowJo (v10.10.1). Cell populations were identified by sequential gating after excluding debris, doublets and dead cells. Representative gating strategies for core panels are provided in Supplementary Figures 1 and 2.

### Antibody-based in vivo perturbations

All cell depletion and cytokine neutralization experiments were performed by intraperitoneal administration of monoclonal antibodies. To prevent CD8+ Trm cell establishment during primary challenge, mice received 0.2 mg anti-CD8αantibody (BioXCell BE0117) on D-1, D2, D5 and D8 of primary MC903 challenge. To deplete circulating CD8+ T cells in recall, mice received 0.2 mg anti-CD8α for three consecutive days before secondary MC903 challenge.

For IFNγ neutralization, mice received 0.2 mg anti-IFNγ antibody (BioLegend 505711) either during primary MC903 challenge (D-1, D2, D5 and D8) or secondary challenge (D35 and D38). For IL-15 neutralization, mice received a single 0.25 mg dose of anti-IL-15 antibody (BioXCell BE0315) one day before secondary MC903. TNF-α neutralization was performed by administering a single 0.5 mg dose of anti-TNFα antibody (BioLegend 506348) one day before secondary MC903 challenge.

### Modulation of fibroblast *STAT1* signaling

#### Evaluating STAT1 variants in 3T3 cells

We used two *STAT1* variants. STAT1dn, which corresponds to human Y701F, is a dominant-negative allele that abolishes the activating tyrosine phosphorylation required for *STAT1* activation ^3^. STAT1ca, which corresponds to human A656C/N658C, is an engineered C-terminal double cysteine mutant that forms an inter-monomer disulfide bond to stabilize the active dimerized form ^4,5^.

NIH/3T3 cells were co-transfected with a piggyBac donor transposon and a hyperactive piggyBac transposase to generate stable, non-clonal cell pools ^6^. Each donor plasmid encoded a doxycycline-inducible, mouse codon-optimized *STAT1*-variant open reading frame (RefSeq NP_001192242.1) and a constitutively expressed puromycin-resistance cassette. The PB-TRE-EGFP-EF1a-rtTA backbone was a gift from Volker Busskamp (Addgene 104454). Donor plasmid was combined with a plasmid encoding the hyperactive piggyBac transposase and delivered using Lipo-fectamine 3000 (ThermoFisher L3000015) according to the manufacturer’s protocol. Following transfection, transposon-integrated cells were selected and expanded in 2 µg/mL puromycin, which was maintained throughout the experiment.

To induce *STAT1*-variant expression, cells were treated with doxycycline (10 µg/mL) or vehicle (0.1% DMSO) for 72 hours. Cells were then stimulated with 10 ng/mL recombinant mouse IFN-γ (BioLegend 575304) for 0 or 24 hours in triplicate. Cells were lysed in Buffer RLT containing 1% β-mercaptoethanol for bulk RNA-sequencing.

#### AAV-based modulation of STAT1 signaling

The pAAV-CAG-GFP plasmid was a gift from Edward Boyden (Addgene 37825). *STAT1* variants were cloned with an N-terminal FLAG tag into this backbone. Plasmids were packaged into AAV-PHP.S vectors and purified by iodixanol density gradient either in-house following established protocols or commercially by Welgen ^7^. For pilot transduction experiments, viral genome titers were determined by qPCR against the internal WPRE in the viral genome. 20 µL of AAV was injected intradermally into the dorsal side of the ear skin at a concentration of 1-5 × 10^11^ genomic copies/mL three days before secondary MC903 challenge. Transduction efficiency was assessed with flow cytometric quantification of intracellular FLAG staining or a fluorescent reporter, as appropriate.

### Pharmacologic CXCR4 inhibition

CXCR4 signaling was inhibited using the peptide antagonist EPI-X4 (MedChemExpress HY-P10415) ^8^. Mice received 400 nmol EPI-X4 in 20 µL of 80% ethanol to the left MC903-treated ear (10 µL each to the dorsal and ventral surfaces) on alternating days of primary MC903 challenge (D1, D3, D5, D7, D9 and D11). Contralateral right ears received 20 µL of 80% ethanol vehicle alone. On days where both MC903 and Epi-X4 were administered (D1 and D5), EPI-X4 was applied 6-8 hours following MC903 treatment.

### Ear skin explant assays

For ex vivo rechallenge experiments, mice were subjected to primary MC903 challenge and euthanized after recovery on D36. Immediately following euthanasia, MC903 (2 nmol in 20 µL) was applied to both the previously challenged left ear and the contralateral vehicle-treated right ear. After allowing MC903 to dry for approximately 1 minute, ears were harvested, separated into dorsal and ventral halves, and floated dermis-side down in 1.5 mL D10 medium (DMEM, 10% FBS, 10 mM HEPES, 2 mM GlutaMAX, 0.1 mM NEAA and 1 Mm sodium pyruvate). Explants were then cultured for 20 hours at 37°C and 5% CO2.

Following explant culture, tissues were homogenized in Buffer RLT (Qiagen 79216) containing 2% β-mercaptoethanol using Lysing Matrix D tubes (MP Biomedicals) and a Bead Ruptor homogenizer at 4°C with 3 cycles at 6.0 m/s and a 5-minute rest between cycles. Homogenates were clarified by centrifugation at 14,000× g for 10 minutes, and total RNA was isolated using the RNeasy Mini Kit (Qiagen 74106) according to the manufacturer’s instructions. RNA was treated with DNase I (ThermoFisher EN0521) and reverse-transcribed using the RevertAid Reverse Transcription Kit (ThermoFisher K1691). Mouse *Ifng, Cxcl9, Cxcl10*, and *Ccl19* transcripts were then quantified by qPCR using TaqMan Gene Expression Assays.

### Primary dermal fibroblast cultures

#### Primary dermal fibroblast isolation

To establish primary dermal fibroblast cultures, ears were separated into dorsal and ventral halves and floated dermisside down in 5 mg/mL Dispase II for 45 minutes at 37°C with gentle agitation. Tissue was then minced and digested in DMEM supplemented with 10 mM HEPES, 250 µg/mL Liberase TM and 250 µg/mL DNase I for 1 hour at 37°C with agitation. Liberated cells were filtered through 70-µm cell strainers, subjected to ACK Lysing Buffer, and plated in D10 supplemented with antibiotic-antimycotic. After 48 hours, culture medium was replaced to remove non-adherent cells, and fibroblasts were expanded to confluence. Following initial plating, fibroblasts were passaged two to three times for expansion before downstream experiments.

#### Quantitative PCR of stimulated fibroblasts

Before cytokine stimulation, fibroblasts were serumstarved overnight. Cells were then stimulated for 48 hours with TNF-α or IL-4 at a final concentration of 20 ng/mL. Cells were lysed in Buffer RLT containing 1% β-mercaptoethanol, triturated, and frozen on dry ice. cDNA was subsequently generated using the reverse transcription portion of the SMART-seq2 (SS2) protocol as previously described ^9^. *Ccl19* and *Tnfaip3* transcripts were then quantified using TaqMan Gene Expression Assays.

### Bulk RNA-seq of fibroblast cultures

For bulk RNA sequencing experiments, primary dermal fibroblasts were isolated from Pdgfra-eGFP reporter mice. Fibroblasts isolated from both ears of each mouse were pooled to generate a single biological replicate. Following initial plating, cells were harvested at passage 0, stained with Zombie NIR Fixable Viability Dye (BioLegend), and live eGFP+ fibroblasts were sorted using a BD FACSMelody cell sorter to minimize contamination by non-fibroblast cell populations. Before cytokine stimulation, cultures were serumstarved overnight and stimulated with individual or combinatorial cytokine treatments consisting of IL-4, IL-13, IFNγ and TNF-α for 24 hours. Each cytokine was used at a final concentration of 10 ng/mL. Cells were lysed in Buffer RLT containing 1% β-mercaptoethanol, and cDNA libraries were generated using the complete SS2 protocol as previously described ^9^. Libraries were sequenced to a target depth of 5 million reads per sample on an Illumina sequencer of appropriate scale. Raw BCL files were demultiplexed using bcl2fastq and aligned to the GRCm39 mouse reference genome using HISAT2^10^.

### CD8+ T cell migration assays

CD8+ T cells were isolated by mechanically dissociating spleens through 70-µm cell strainers to generate singlecell suspensions, followed by ACK Lysing Buffer treatment. CD8+ T cells were then purified by negative selection using the Mouse CD8+ T Cell Isolation Kit (Miltenyi 130-104-075). Purified CD8+ T cells were cultured in RP10 medium (RPMI1640, 10% FBS, 10 mM HEPES, 2 mM GlutaMAX, 0.1 mM NEAA, 1 mM sodium pyruvate, and 1 × antibiotic-antimycotic) and activated with Dynabeads Mouse T-Activator (ThermoFisher) for 48 hours according to the manufacturer’s instructions. Dynabeads were magnetically separated, and activated CD8+ T cells were cultured for an additional 24 hours before use in migration assays.

Explant-conditioned medium was collected from ex vivo explant recall assays. Migration assays were performed using 24-well Transwell inserts with 5-µm pore membranes (Corning). Activated CD8+ T cells (5 × 10^5^ cells) were suspended in 160 µL D10 medium and added to the apical chamber, while 600 µL explant-conditioned medium was added to the basal chamber. Cells were allowed to migrate for 3 hours at 37°C and 5% CO_2_. Migrated and unmigrated cells were subsequently collected, stained with Zombie NIR Fixable Viability Dye and Alexa Fluor 488-conjugated anti-CD8α antibody, and quantified by flow cytometry using 123count eBeads to determine the ratio of migrated to unmigrated CD8+ T cells.

### Longitudinal whole-skin scRNA-seq

Longitudinal whole-skin scRNA-seq was performed across primary and secondary challenge for both MC903 and IMQ. Three biological replicates, each consisting of a single mouse, were collected at every timepoint. Ear skin was harvested immediately following euthanasia and washed extensively in HBSS before manual separation into dorsal and ventral halves. Epidermis and dermis were separated by incubation in 5 mg/mL Dispase II for 40 minutes at 37°C with agitation. Dermal tissue was minced and digested in DMEM containing 100 µg/mL Liberase TM, 100 µg/mL DNase I, 1.5 mg/mL hyaluronidase, 10 mM HEPES and 2% FBS for 1 hour at 37°C with agitation. Epidermal sheets were minced and dissociated in 0.25% trypsin-EDTA for 10 minutes at 37°C with agitation. Dermal and epidermal cell suspensions were independently filtered through sequential 70-µm and 40-µm cell strainers, subjected to ACK Lysing Buffer, and counted before downstream processing.

For each biological replicate, matched dermal and epidermal cell suspensions were recombined at a 4:1 ratio prior to hashing to ensure adequate dermal representation given the expected epidermal hyperplasia in these inflammatory models ^2,11^. Cells were blocked with TruStain FcX PLUS before staining with a cocktail of biotinylated antibodies against EpCAM, PDPN, CD31, H-2K, CD45, CD44, Thy1.2, PDGFRβ and PDGFRα for 30 minutes at 4°C as previously described ^12^. After washing, cells were incubated with TotalSeq-B Streptavidin-B0951-0955 (BioLegend) for 20 minutes at 4°C with each biological replicate labeled using a unique barcode. Hashed biological replicates were counted and pooled at a 1:1:1 ratio corresponding to each experimental timepoint. Single-cell gene expression libraries were generated using the Chromium Next GEM Single Cell 3’ (v3.1 Chemistry Dual Index) with Feature Barcoding technology for Cell Surface Proteins (10x Genomics). Gene expression and hash libraries were pooled at a 15:1 ratio and sequenced on the appropriate Illumina platform to achieve a targeted depth of 30,000 reads per cell.

### scRNA-seq of CD8+ T Cells

CD8+ T cells were profiled from recovered D36 skin and D37 skin after one day of MC903 recall. Ten biological replicates were profiled at each timepoint, with one mouse comprising each biological replicate. Ear skin was harvested immediately after euthanasia, separated into dorsal and ventral halves, and digested according to the immune cell digestion protocol. Single-cell suspensions were stained with Zombie Violet, blocked with TruStain FcX PLUS prior to staining for 30 minutes at 4°C with biotinylated anti-CD45, PE/Cy7-conjugated anti-CD8β, and TotalSeqC oligonucleotide-conjugated antibodies against CD103, CD69, CD62L and CD44. After washing, cells were incubated with TotalSeq-C PE-Streptavidin barcodes (BioLegend) for 20 minutes at 4°C. Pooled samples were washed, filtered and sorted on a BD FACSMelody cell sorter to isolate Live CD45+ CD8β+ cells.

Sorted cells were washed, resuspended in PBS with 1% BSA, and loaded onto the Chromium GEM-X Single Cell 5′ v3 Gene Expression with Feature Barcoding technology for Cell Surface Protein (10x Genomics). Gene expression, TCR and Feature Barcode (CITE-seq) libraries were pooled at a 10:1:1 ratio and sequenced on an Illumina sequencer targeting 30,000 reads per cell.

### scRNA-seq preprocessing and annotation

scRNA-seq data were demultiplexed and aligned to the GRCm39-2024-A mouse reference genome using CellRanger (v9.0.1). Raw BCL files were demultiplexed using bcl2fastq and the cellranger multi function was used to call the appropriate count function for gene expression, cell surface protein and TCR libraries. Ambient RNA background was corrected using cellbender (v0.3.0) ^13^. Downstream analysis was conducted using Seurat (v5.1.0) ^14^.

All samples were merged into one object and initial quality control filtering was conducted using Unique Molecular Identifier count (< 40000), number of genes detected (< 7000) and percentage of mitochondrial reads (< 15%). Cells were normalized using scTransform (v0.4.1), followed by PCA, Louvain clustering and UMAP embedding. To identify cell states, iterative clustering was performed as previously described ^15^. Each broad cluster was separated for secondary filtering by quality control metrics, re-clustered and annotated. At each level of annotation, doublets were identified based on coexpression of multiple lineage markers. Hash counts were normalized and assigned to individual mice using the HTOdemux function in Seurat.

### scRNA-seq compositional analysis

Compositional analysis was performed using only cells confidently assigned as a singlet by HTOdemux. After annotation of all end cell clusters, cell state abundances were calculated by dividing the number of cells belonging to a state by the total number of cells for that mouse replicate. A pseudocount was generated by multiplying each frequency by 3000, as previously described ^16^. Pseudocounts were transformed using a centered log-ratio (CLR) transformation, which was the abundance matrix used for compositional PCA analysis as previously described ^16^. This abundance matrix was also used as input for proportionality analysis performed using the propr package (v5.0.0) ^17^. A significance cutoff FDR < 0.05 generated by permutation testing was used to identify significant relationships between cell clusters. Cellcell communication was inferred using CellChat (v2.1.0) with log-normalized counts as input. CellChat nominates ligand-receptor pairs based on differential expression and levels of downstream signaling mediators ^18^.

### scRNA-seq differential expression analysis

Differential gene expression analysis was conducted at the single-cell level for cell state marker annotation. The Seurat FindAllMarkers function was used to perform a Wilcoxon rank-sum test and significant genes were filtered to require expression in at least 20% of cells and a log fold-change of at least 0.25. Marker annotations for end cell clusters are provided in Supplementary Table 1.

Differential gene expression analysis was also conducted at the pseudobulk level. Gene expression was aggregated across individual mice and cell clusters where appropriate using the Seurat AggregateExpression function. Pseudobulked samples were normalized and differential gene expression was tested using DESeq2 (v1.52.1) ^19^. Gene set enrichment analysis was conducted using fgsea (v1.32.2) restricted to hallmark gene sets^20^.

### Integrated atopic dermatitis scRNA-seq dataset generation

An integrated human AD atlas was generated from five publicly available studies comprising healthy control, AD non-lesional and AD lesional skin samples. Datasets included He et al, JACI 2020 (GSE147424), Rojahn et al, JACI 2020 (GSE153760), Bangert et al, Sci Immunol 2021 (GSE158432), Bangert et al, Nat Commun 2024 (GSE230575) and Fiskin et al, Nat Commun 2026 (GSE204765) ^21–25^. Only healthy control, AD non-lesional and AD lesional skin samples from each study were included; all other sample types and conditions were excluded.

Datasets were integrated using Harmony to correct for batch effects using study ID and patient ID as batch variables^26^. Major cell types were manually annotated based on canonical marker gene expression (Keratinocyte: *KRT5, KRT14, KRT1, KRT10*; Fibroblast: *PDGFRA, COL1A1, COL1A2*; Myofibroblast: *ACTA2, TAGLN*; Glial Cell: *PLP1, SOX10, S100B, MPZ*; Melanocyte: *MLANA, TYR, DCT*; Vascular Endothelium: *PECAM1, FLT1, SELE*; Lymphatic Endothelium: *PECAM1, PROX1, CCL21*; Lymphocyte: *CD3E, CD3D*; Myeloid Cell: *HLA-DRA, CD68*). Fibroblast and T cell populations were subsequently extracted for independent analysis. Within the fibroblast and T cell objects, doublets were identified and removed before iterative clustering and manual annotation of transcriptionally distinct states. Cell states were assigned based on a combination of canonical lineage markers and differentially expressed genes.

### Analysis of integrated AD scRNA-seq dataset

#### Disease severity association analysis

Associations between fibroblast or T cell state abundance and disease severity were assessed using Validated Investigator Global Assessment for Atopic Dermatitis (vIGA-AD) metadata where available. For each patient, the proportion of cells occupying each annotated fibroblast or T cell state was calculated relative to the total number of cells within that lineage. Associations between patient-level cell state abundance and vIGA-AD score were assessed using Spearman’s rank correlation. Spearman’s correlation coefficient (ρ) and corresponding p-value were calculated for each annotated cell state using Benjamini-Hochberg correction.

#### Pseudobulk differential expression analysis of AD Trm

Pseudobulk differential expression analysis was performed on AD Trm states where matched treatment metadata was available. Raw counts were aggregated for each patient-treatment-state combination using Seurat v5 AggregateExpression, summing counts across all cells within each pseudobulk sample. Only patient-state combinations containing at least 10 cells were retained for downstream analysis. Differential gene expression between untreated lesional AD skin and dupilumab-treated post-lesional skin was performed using DESeq2^19^, with samples collected after 16 weeks and 1 year of dupilumab treatment combined into a single treated cohort.

#### Cross-species gene signature mapping and gene set variation analysis

Cross-species comparison of allergic CD8+ Trm cells and inflammatory fibroblasts was performed by generating murine gene signatures from the corresponding mouse scRNA-seq datasets. The allergic CD8+ Trm signature was defined as the top 100 differentially expressed genes defining the D37associated *Ccl1*+ CD8+ Trm state ranked by adjusted p-value. The inflammatory fibroblast signature was defined from the longitudinal whole-skin scRNA-seq dataset and consisted of the top 50 differentially expressed genes between D37 and D36 fibroblasts ranked by adjusted p-value. Murine gene signatures were converted to human orthologs using the orthogene package (v1.18.1) with the babelgene method, retaining only one-to-one orthologs (non121_strategy = 3) ^27^.

Gene module activity was quantified in pseudobulk patientstate profiles using gene set variation analysis (GSVA) (v2.6.5) ^28^. Briefly, pseudobulk expression matrices were generated using Seurat v5 AggregateExpression after filtering patient-state combinations containing fewer than 10 cells. GSVA scores were calculated using the GSVA package with Gaussian kernel density estimation (kcdf = “Gaussian”) and a minimum gene set size of 10 genes^28^. Gene module activity was scored independently for murine allergic CD8+ Trm cell and inflammatory fibroblast signatures using their corresponding human ortholog gene sets.

### Statistical Testing

Exact replicates for each experiment are listed in figure legends. All experiments were successfully repeated at least three times. Statistical methods to predetermine sample size were not used. All data are represented as mean*±* s.e.m. Comparisons between groups were performed using unpaired two-tailed t-tests or one-way analysis of variance with post-hoc Tukey’s multiple comparison unless otherwise stated. Across all statistical tests, p < 0.05 was considered significant. Across all plots, statistical significance is denoted as follows: * p *≤* 0.05; ** p *≤* 0.01; *** p *≤* 0.001; and **** p *≤* 0.0001. All statistical tests were conducted in R (v4.5.1) or GraphPad Prism (v10.6.1).

## Data Availability

scRNA-seq data will be made available for download and visualization via the Broad Institute Single Cell Portal (https://singlecell.broadinstitute.org/single_cell) upon publication. RNA-seq and scRNA-seq datasets will be deposited to the Gene Expression Omnibus.

## Code Availability

Code was written in R version 4.5.1 using publicly available packages. No new algorithms or functions were created and all code relied on built-in functions in publicly available packages on CRAN and Bioconductor. Code will be made available on GitHub in the jo-m-lab repository (https://github.com/jo-m-lab) upon publication.

## Extended Data Figures

**Extended Data Figure 1.**
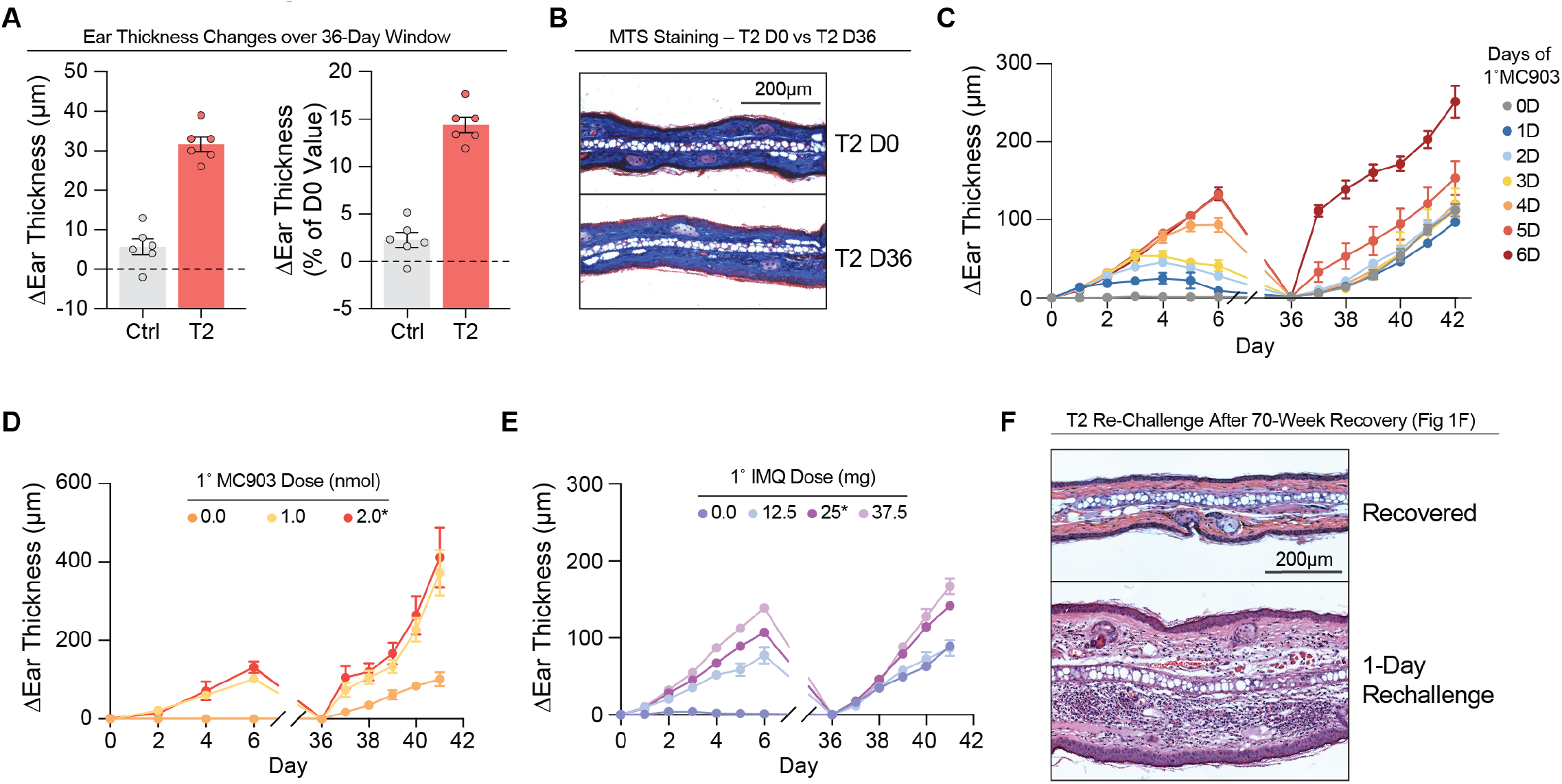
Characterizing properties of T2 and T17 inflammatory memory. **(A)** Change in ear thickness between D36 and D0 for vehicle-treated control right ear and MC903-treated (T2) left ear displayed as absolute value and percent change from starting ear thickness (n=6 mice). **(B)** Representative image of Masson’s trichrome stain (MTS) of D0 and T2-treated D36 ears, where blue represents collagen deposition. **(C)** Ear thickness curve for allergic memory recall after different lengths of primary T2 challenge (n=3 mice/group). **(D)** Ear thickness curve for allergic memory recall with 0, 1 or 2 nmol MC903 used in primary challenge (n=4 mice/group). Asterisk denotes MC903 dosage used throughout this study. **(E)** Ear thickness curve for T17 memory recall with 0, 12.5, 25 or 37.5 mg IMQ used in primary challenge (n=4 mice/group). Asterisk denotes IMQ dosage used throughout this study. **(F)** Representative H&E image of one-day T2 recall response after prolonged 70-week recovery period.

**Extended Data Figure 2.**
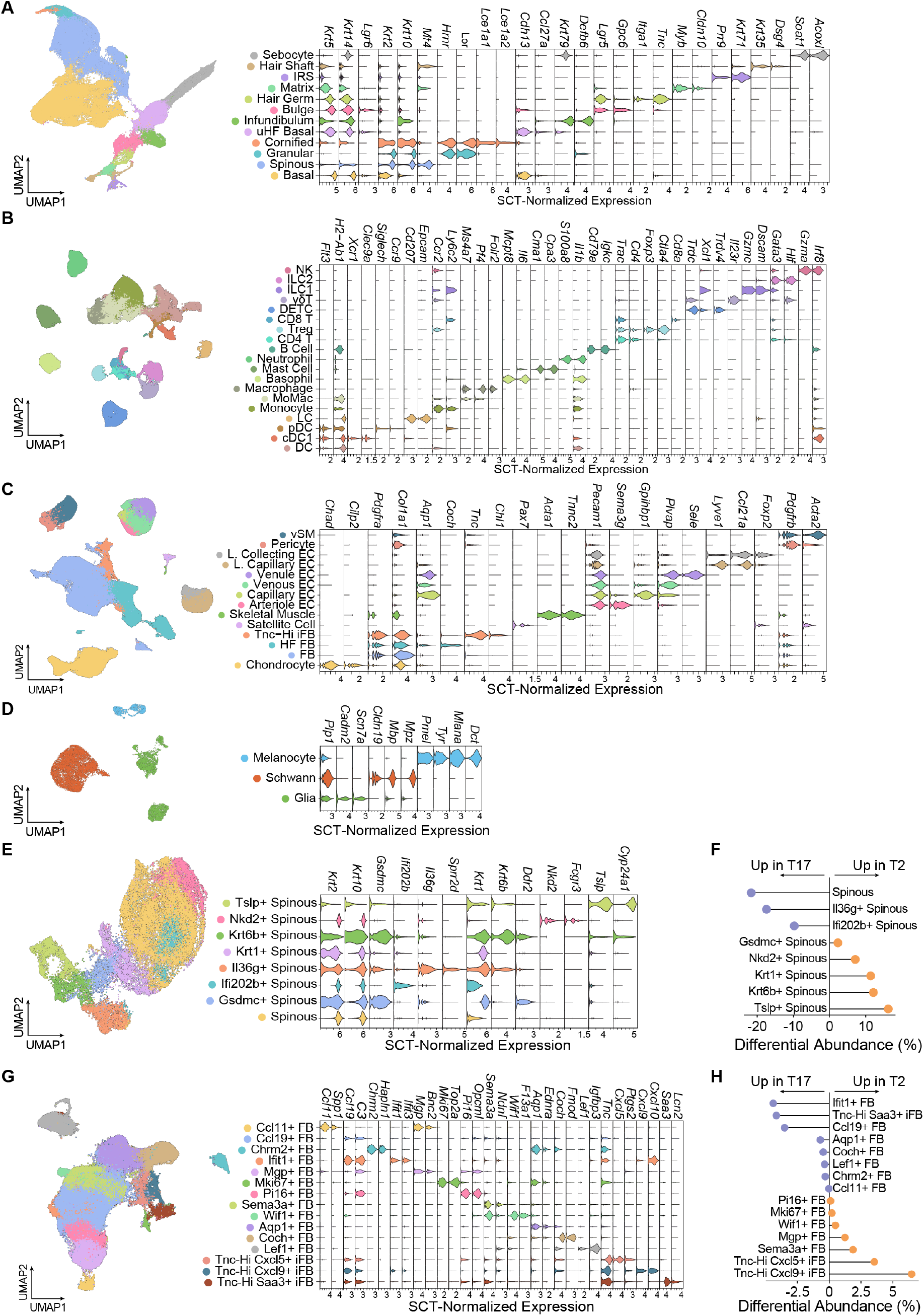
Longitudinal scRNA-seq atlas of murine T17 and T2 skin inflammation. **(A)** UMAP embedding of 84,488 epithelial cells across 12 cell types with representative markers. **(B)** UMAP embedding of 55,654 immune cells across 19 clusters with representative markers. **(C)** UMAP embedding of 131,035 stromal cells across 14 cell types and subtypes with representative markers. **(D)** UMAP embedding of 16,712 neural crest cells across 3 cell types with representative markers. **(E)** UMAP embedding of 26,958 spinous keratinocytes across 8 cell clusters and representative markers for each cluster. **(F)** Enrichment of individual spinous keratinocyte clusters per inflammatory Type was calculated as the differential abundance of the given cluster across all T2 and T17 timepoints. **(G)** UMAP embedding of 34,034 fibroblasts across 15 cell clusters and representative markers for each cluster. **(H)** Enrichment of individual fibroblast clusters per inflammatory Type was calculated as the differential abundance of the given cluster across all timepoints of T2 or T17 inflammation.

**Extended Data Figure 3.**
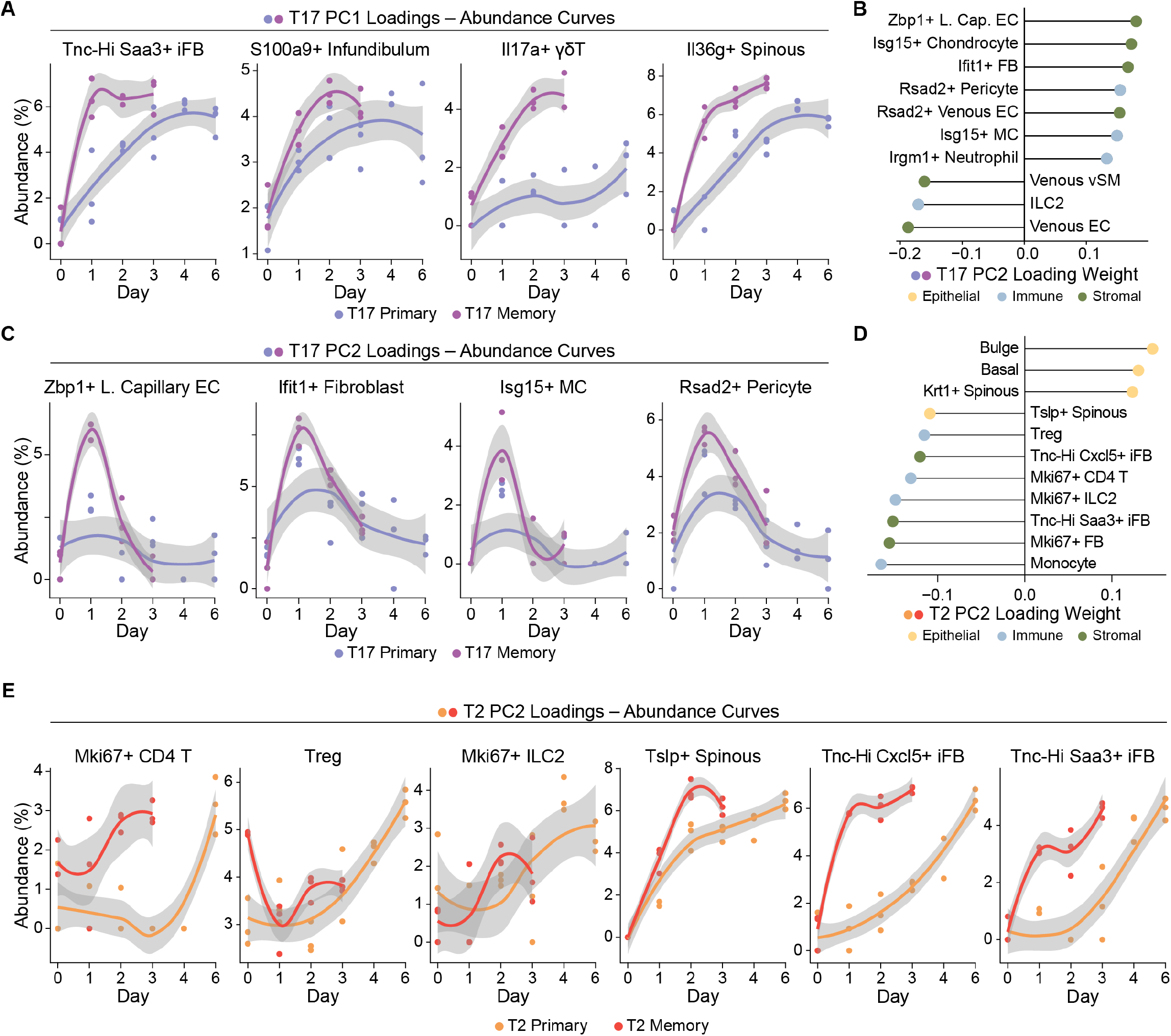
Compositional analysis of T17 and T2 skin inflammation. **(A)** Longitudinal abundance curves, calculated as representation within the total tissue ecosystem, of PC1 loadings for T17 compositional PCA from Fig 2C. **(B)** PC2 loadings for T17 compositional PCA and **(C)** representative abundance curves relative to the total tissue ecosystem. **(D)** PC2 loadings for T2 compositional PCA (Fig 2G) and **(E)** representative abundance curves relative to the total tissue ecosystem.

**Extended Data Figure 4.**
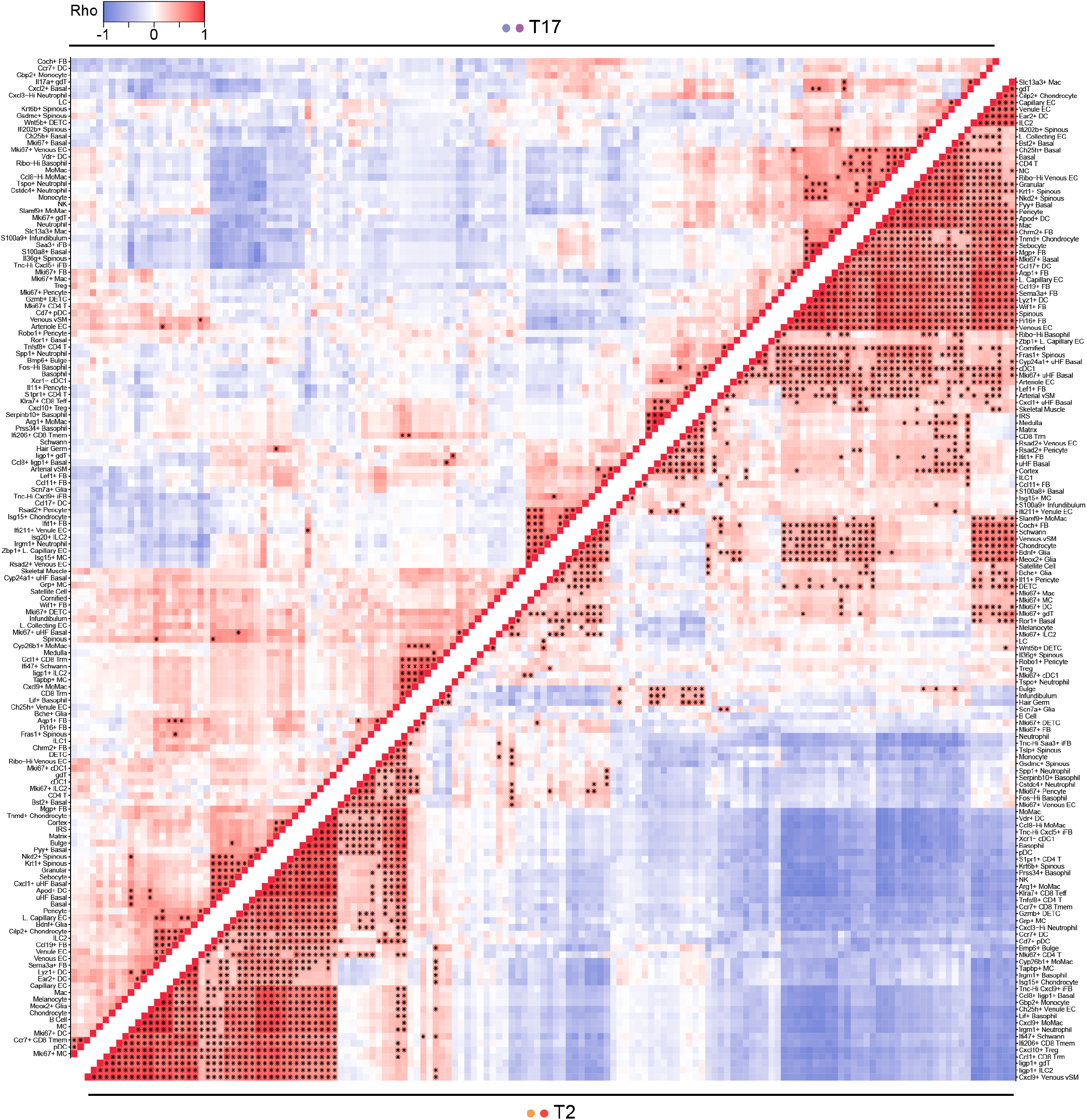
Proportionality networks for T17 and T2 inflammatory scRNA-seq timecourses. Hierarchically clustered similarity heatmap of mouse replicate proportionality measured by Rho coefficient calculated across all timepoints for T17 (top) and T2 (bottom) inflammation. Significant correlations between pairs of clusters are depicted with an asterisk calculated by permutation testing with FDR *<* 0.05.

**Extended Data Figure 5.**
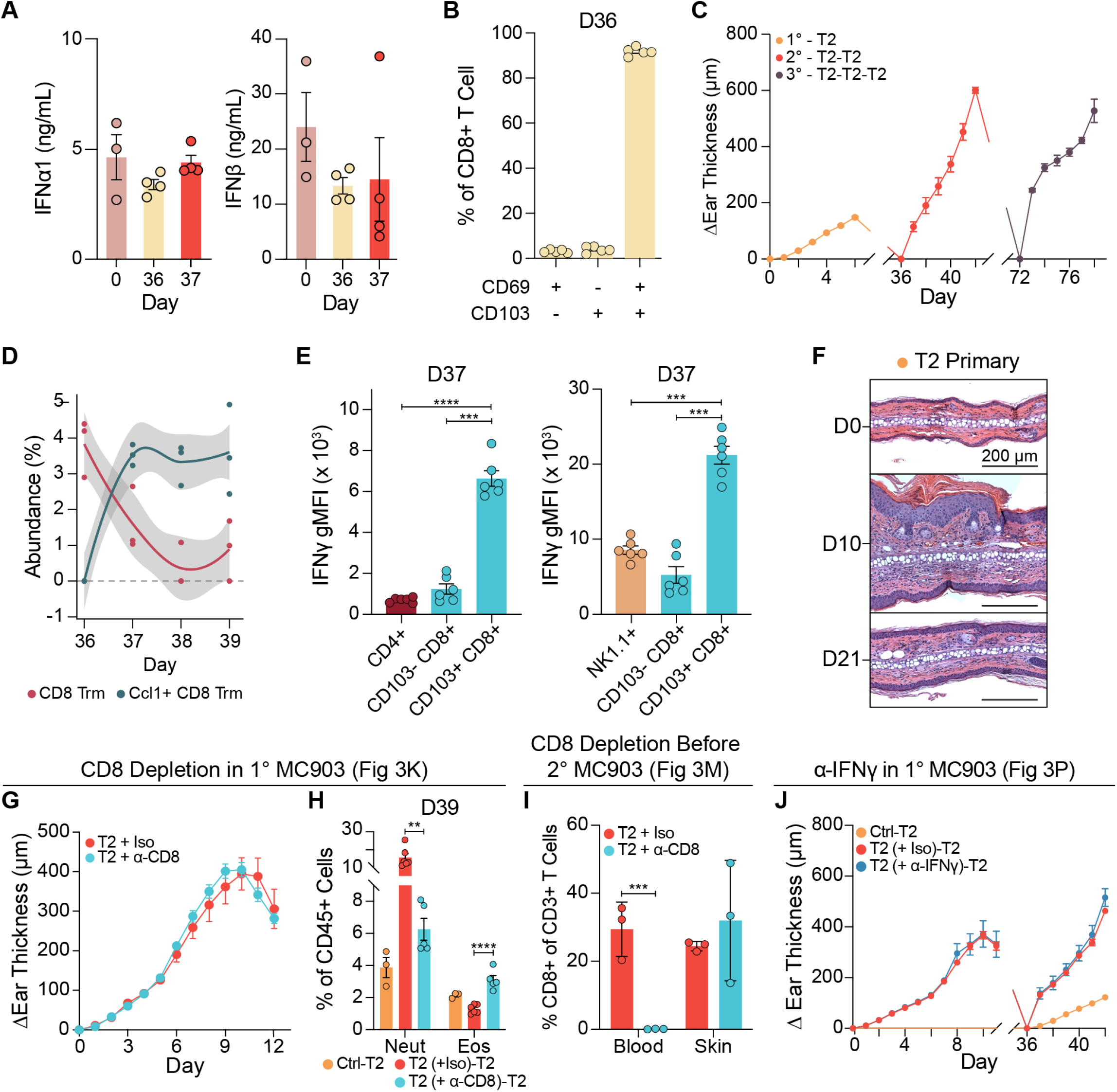
Characterizing the role of CD8+ T cells and IFNγ across acute and memory T2 inflammation. **(A)** Concentration of IFNα1 and IFNβ in ear skin lysates in T2 memory (n=3 (D0), 4 (D36, D37) mice). **(B)** Flow cytometry assessment of residency-associated markers CD69 and CD103 on CD8+ T cells in recovered D36 skin (n=5 mice). **(C)** Ear thickness curves across primary to tertiary MC903 challenge (n=5 mice). **(D)** Abundance of CD8+ Trm and *Ccl1*+ CD8+ Trm relative to full tissue ecosystem over T2 memory timepoints. **(E)** gMFI of IFNγ in D37 skin samples assessed by flow cytometry across 2 different panels to separately capture NK1.1+ and CD4+ T cells (n=6 mice/panel). **(F)** Representative H&E images from later primary T2 timepoints capturing peak (D10) and late resolution (D21). **(G)** Extended primary ear thickness curve during primary CD8+ T cell depletion in T2 inflammation (n=5 mice/group) and **(H)** flow cytometry results on D39 (n=3 (Ctrl-T2), 6 (Iso), 5 (α-CD8) mice). **(I)** Flow cytometric analysis of CD8+ T cells in blood and skin on D36 after circulating CD8+ T cell depletion (n=3 mice). **(J)** Extended ear thickness curve for primary and memory T2 inflammation under primary IFNγ neutralization (n=5 (Ctrl-T2, α-IFNγ), 10 (Iso) mice).

**Extended Data Figure 6.**
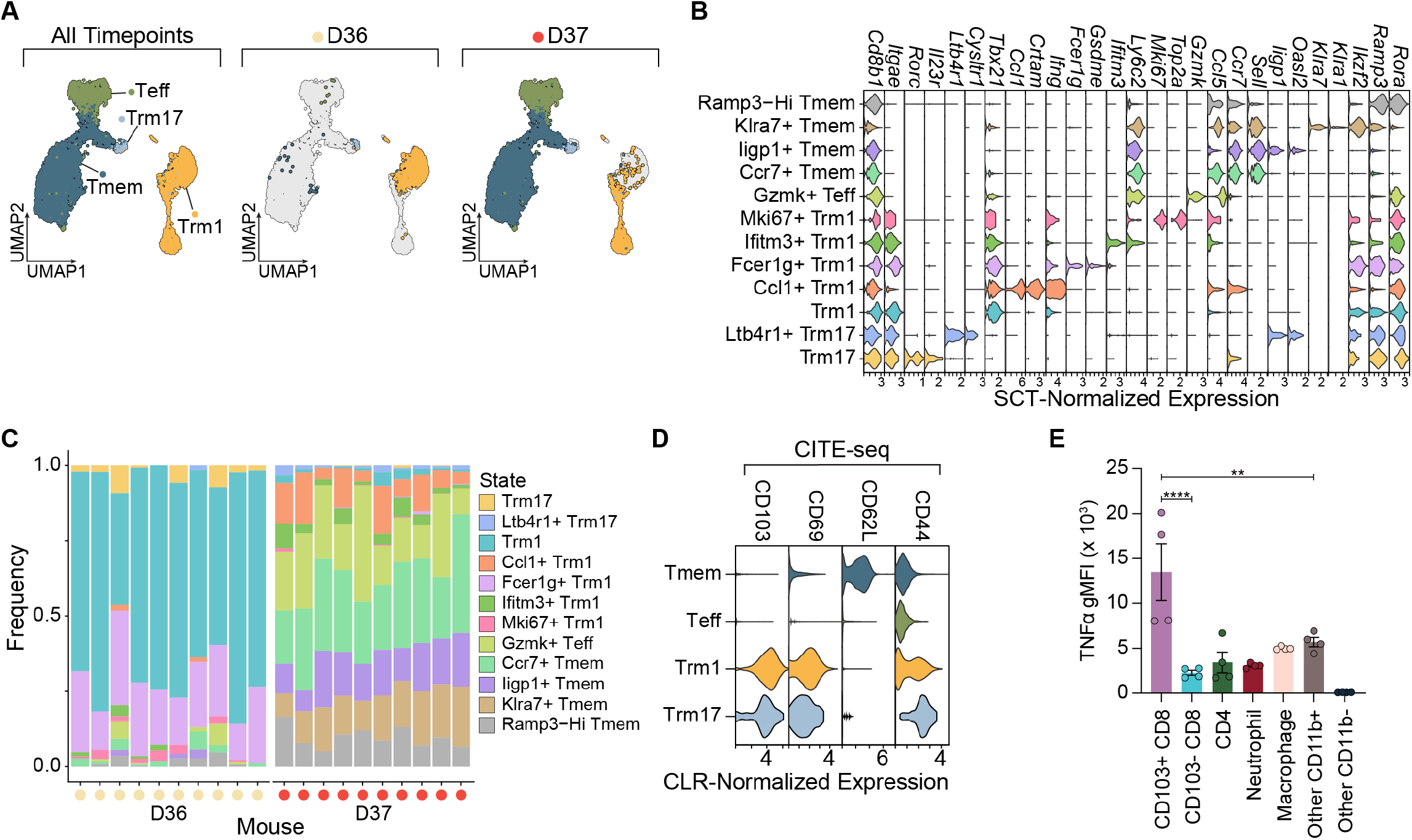
Characterization of murine allergic CD8+ Trm by scRNA-seq and flow cytometry. **(A)** UMAP embedding of 7,708 CD8+ T cells recovered, split by timepoint (n=10 mice/timepoint). **(B)** Violin plot showing representative markers of CD8+ Trm1, Trm17, Teff and Tmem cell states. **(C)** Stacked bar plot showing cell state abundances on a per-mouse basis across 10 mice per timepoint. **(D)** CITE-seq assessment of cell surface CD103, CD69, CD62L and CD44 levels across CD8+ Trm1, Trm17, Tmem and Teff populations. **(E)** Flow cytometric assessment of TNF-α gMFI on D37 across lymphoid and myeloid lineages in skin (n=4 mice).

**Extended Data Figure 7.**
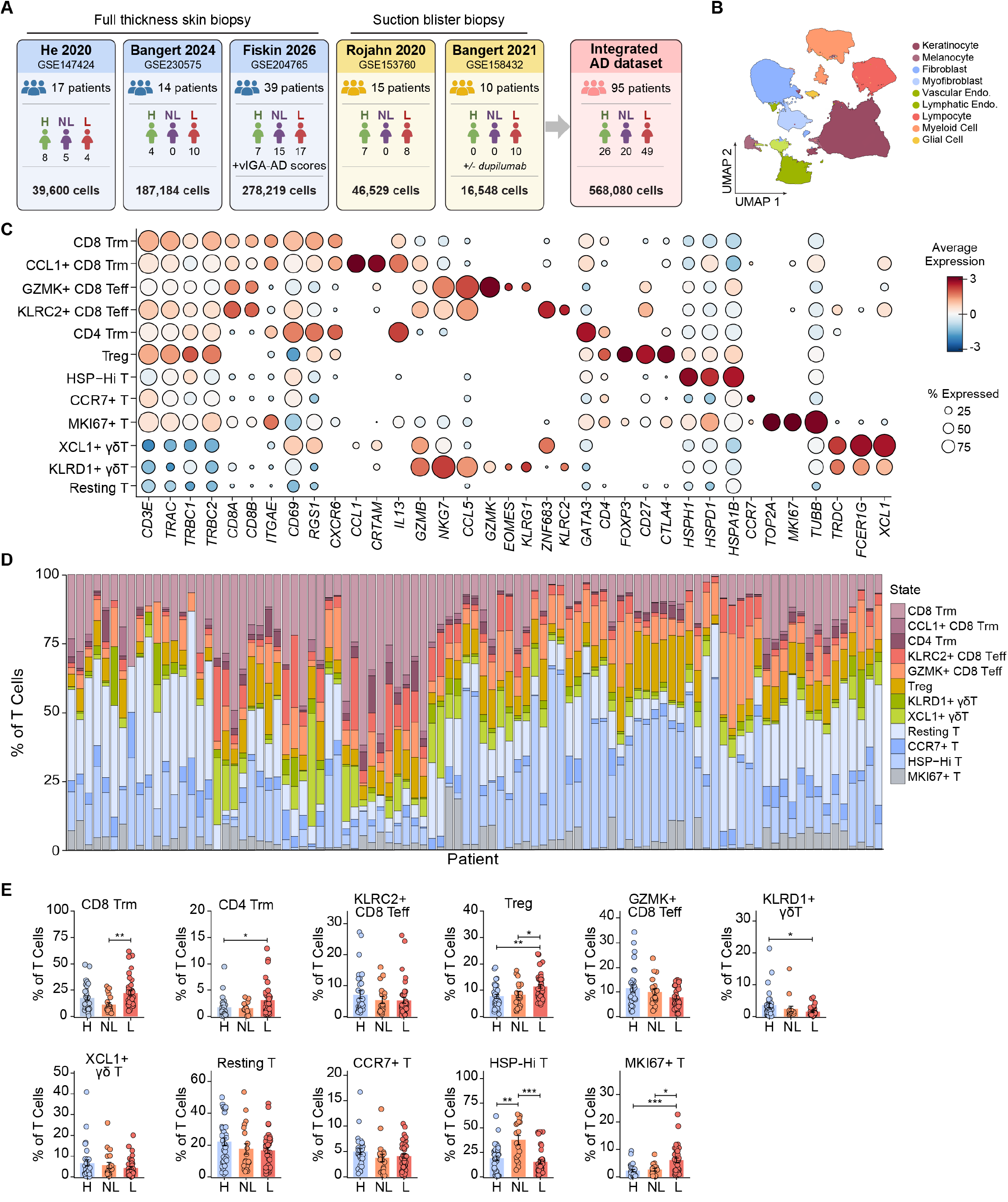
Integrated and re-annotated human T cell atlas in AD. **(A)** 5 public scRNA-seq datasets of AD skin featuring full thickness and suction blister biopsies (GSE147424, GSE230575, GSE204765, GSE153760, GSE158432) were integrated using Harmony and re-annotated to produce a final dataset of 568,080 cells across 95 patients. Additional details in Methods. **(B)** UMAP embedding of 568,080 cells showing annotated cell types from one round of clustering defined using markers specified in Methods. **(C)** T cells (*CD3E*+ *CD3D*+) were subclustered and cluster-defining marker expression is displayed. Related UMAP is shown in Fig 4H. **(D)** Stacked bar chart showing cell patient biopsy. **(E)** Quantification of additional cell clusters in addition to Fig 4K for their abundances in each biopsy across healthy (H), non-lesional (w)oannganesionliavel l (NL) and lesional (L) ski (H),Shift(NL), 41 (L) patients).

**Extended Data Figure 8.**
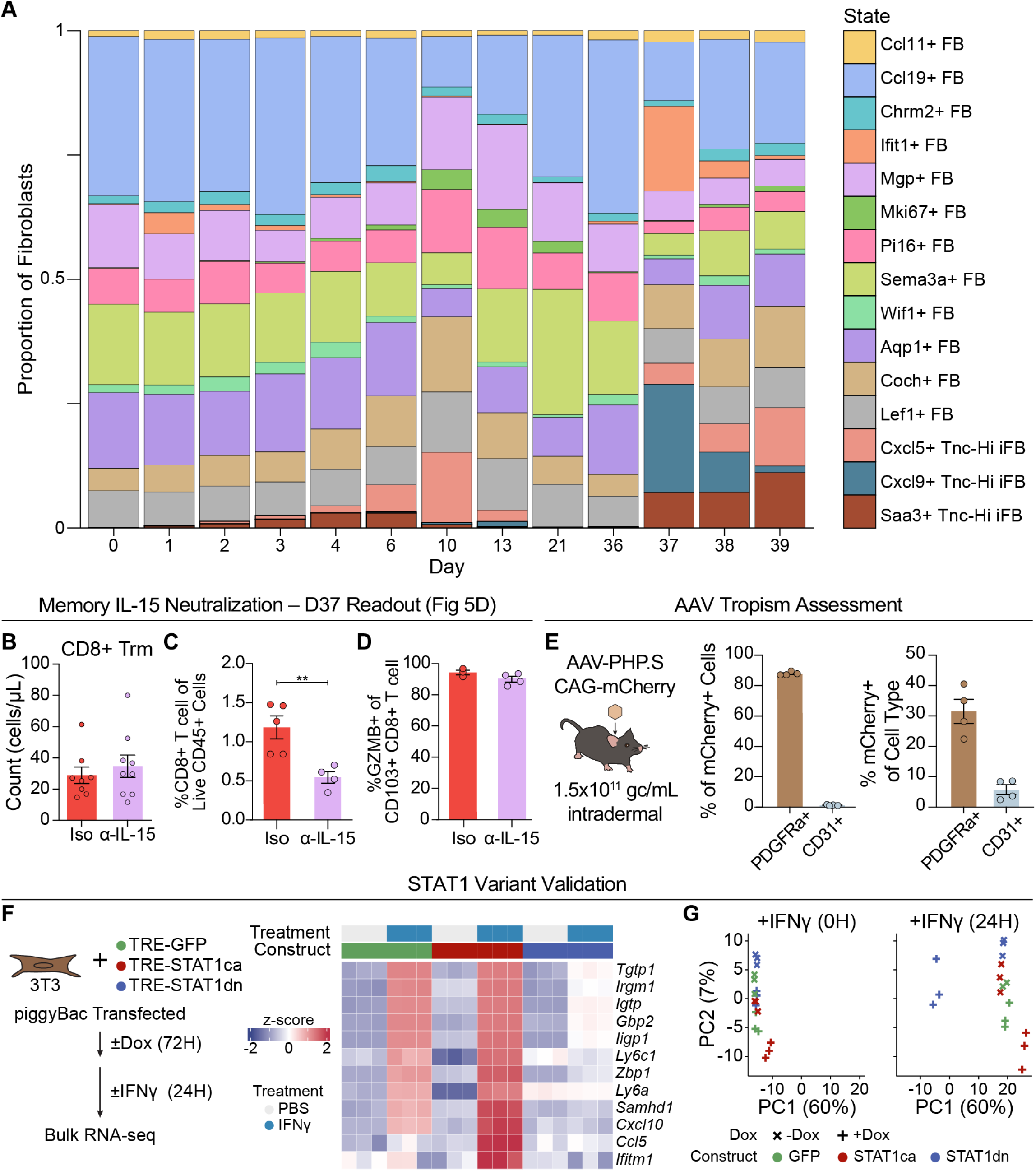
Modulating inflammatory fibroblast activity in murine ear skin. **(A)** Stacked bar plot showing representation of 15 scRNA-seq fibroblast clusters across T2 inflammation timepoints. **(B-D)** Flow cytometric evaluation of **(B)** CD103+ CD8+ Trm counts (n=8 (Iso), 9 (α-IL15) mice), **(C)** CD8+ T cell proportions (n=5 (Iso), 4 (α-IL15) mice), and **(D)** frequency of *GZMB*+ CD8+ Trm (n=3 (Iso), 4 (α-IL15) mice) in skin at D37 after IL-15 neutralization before T2 memory recall. **(E)** Schematic of intradermal AAV injection with CAG-mCherry genome and flow cytometric assessment of PDGFRα+ and CD31+ cell tropism in skin after 3 days (n=4 mice). **(F)** Schematic for generating and stimulating 3T3 stable cell lines with doxycycline-inducible GFP, STAT1ca and STAT1dn plasmids in piggyBac vectors. Heatmap showing significant 24-hour transcriptional changes assessed by bulk RNA-seq (n=3 biological replicates/condition). **(G)** PCA of stably transfected 3T3 cell lines stimulated with IFNγ for 0 or 24 hours with or without 4-hour prior doxycycline induction for expression of *STAT1* variant.

**Extended Data Figure 9.**
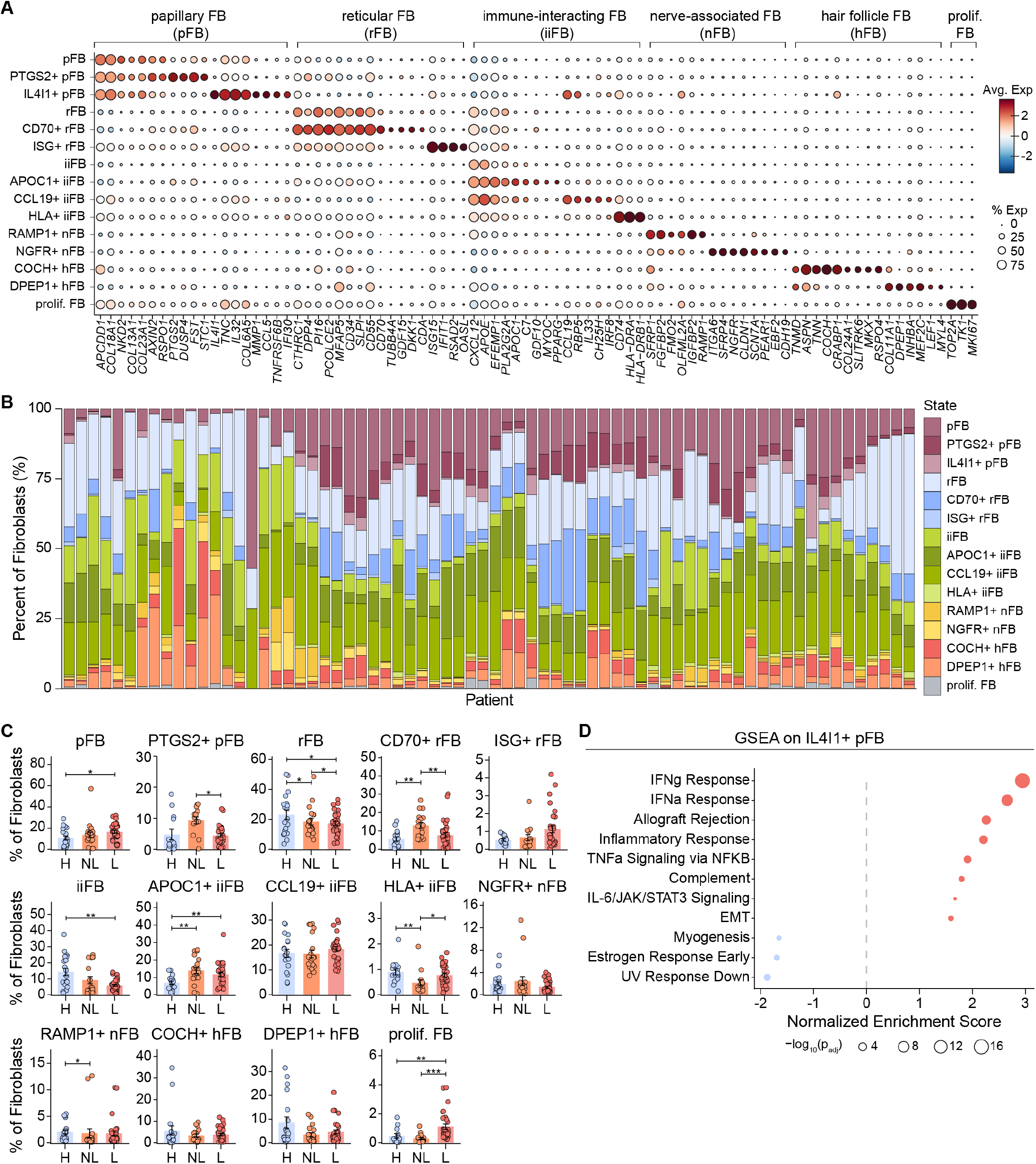
Integrated and re-annotated human fibroblasts in AD atlas. **(A)** Fibroblasts from studies described in Extended Data Fig 7A were clustered to produce associated UMAP in Fig 5L and expression of cluster-defining markers is displayed. **(B)** Stacked bar chart showing cell cluster abundance in each patient biopsy. **(C)** Quantification of fibroblast clusters in addition to *IL4I1*+ pFB in Fig 5O for their abundances in each biopsy across healthy (H), non-lesional (NL) and lesional (L) skin (n=19 (H), 20 (NL), 31 (L) patients). **(D)** State-defining genes for *IL4I1*+ pFB defined at the single-cell level by Wilcoxon rank-sum test were used for GSEA.

**Extended Data Figure 10.**
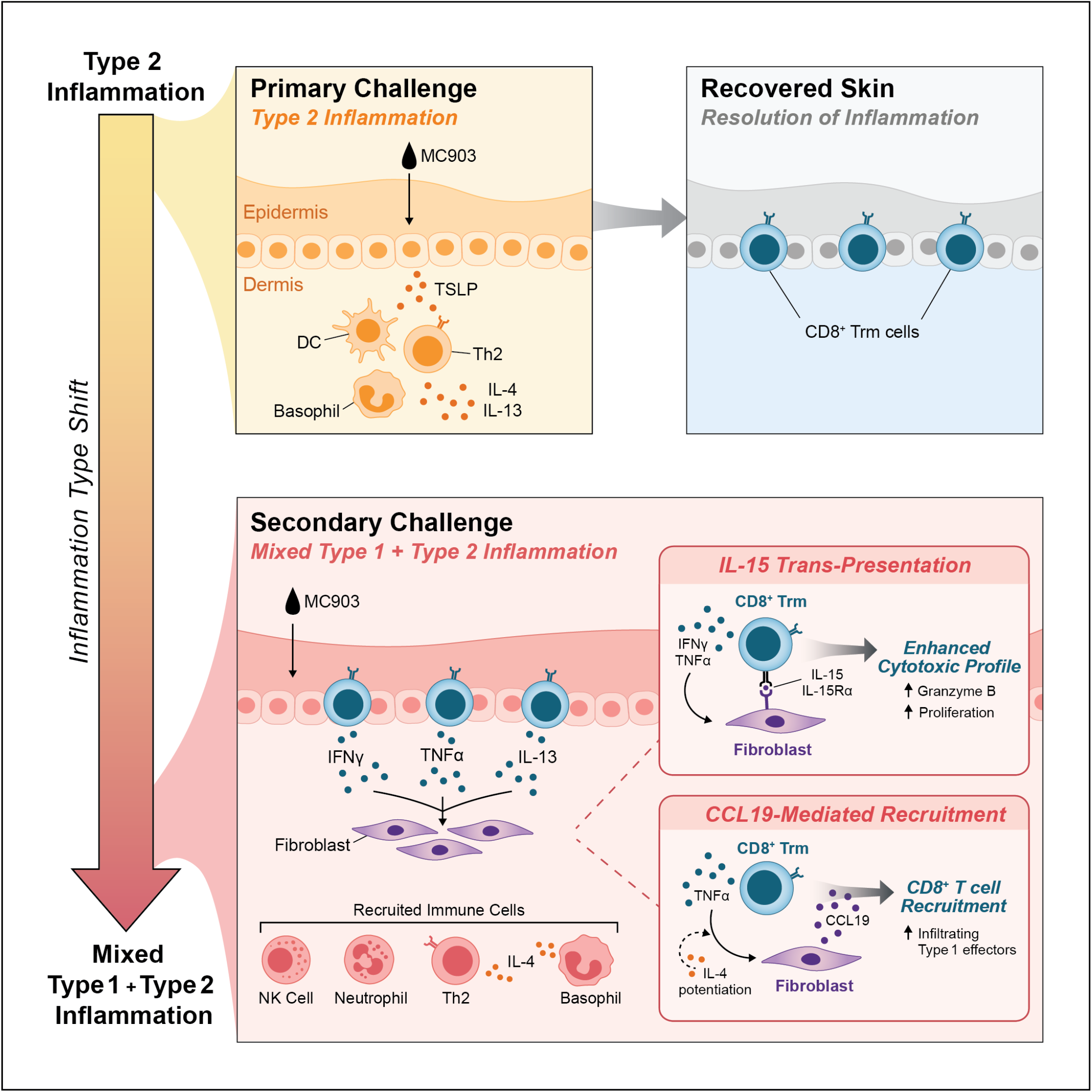
Model of how allergic memory encoding rewires the tissue response to T2 inflammation. Primary allergic inflammation induced by MC903 initiates a TSLP-driven keratinocyte response that promotes a T2 environment driven by IL-4 and IL-13 secretion from Th2 cells and basophils. Upon recovery, epidermal-resident allergic CD8+ Trm cells, a reservoir of inflammatory memory, are established in murine skin. Upon T2 recall, a mixed inflammatory environment is established with rapid recruitment of T1 and T2 effectors. Re-activated allergic CD8+ Trm cells producing IFNγ, TNF-α and IL-13 drive this T2-to-T1 shift in memory and form a two-cell circuit with dermal fibroblasts. Responding to CD8+ Trm-derived T1 effector cytokines, fibroblasts trans-present IL-15 to enhance the effector profile of CD8+ T cells, and recruit circulating T cells via *CCL19* secretion. Concurrent IL-4 signaling potentiates both amplification arms of the fibroblast–CD8+ T cell circuit.

## List of Supplementary Data Files

**Supplementary Table 1: Differentially expressed genes across end cell states called in Fig 2**. Differential expression analysis was performed across all samples within each higher-level classification, denoted by the names of individual sheets, using a Wilcoxon rank-sum test.

**Supplementary Table 2: Differentially expressed genes across end cell states called in Fig 4 for sorted CD8+ T cells**. Differential expression analysis was performed across all samples within each cell subset, denoted by the names of individual sheets, using a Wilcoxon rank-sum test.

**Supplementary Table 3: Flow cytometry panels used across paper**. For each panel, the antibody, clone, vendor and catalog number are provided. Panels have been grouped to describe general immune profiling, targeted Trm cell profiling, cytokine- or chemokine-related profiling, and structural cell-related investigations. Panels from each group are placed on separate sheets.

**Supplementary Figure 1: Representative gating strategies for myeloid and lymphoid panels in skin. (A)** Myeloid cell gating strategy for identification of neutrophils, eosinophils, monocytes, macrophages, dendritic cells and mast cells. **(B)** Lymphocyte gating strategy for identification of CD8+ T cells, CD4+ T cells with further classification as Treg or Th2 cells.

**Supplementary Figure 2: Representative gating strategies for Trm cells and structural cell panels in skin. (A)** Trm cell gating strategy for identification of CD8+ Trm cells as CD103+ CD62L-along with expression of IFNγ and IL-13. **(B)** Structural cell panel for identification of immune, fibroblast and endothelial expression of IL-15Rα.

